# Determinants of hyena participation in risky collective action

**DOI:** 10.1101/2022.05.30.494003

**Authors:** Tracy M. Montgomery, Kenna D.S. Lehmann, Samantha Gregg, Kathleen Keyser, Leah E. McTigue, Jacinta C. Beehner, Kay E. Holekamp

## Abstract

Many species engage in risky cooperative behaviors, which pose a challenge to evolutionary theory: participants take on all the costs of cooperation, yet even non-participants benefit from success in these encounters. So, why participate in these risky behaviors? We address this question using data from spotted hyenas fighting with lions. Lions are much larger, and kill many hyenas, so these fights require cooperative mobbing by hyenas for them to succeed. We identify factors that predict: (1) when hyena groups engage in cooperative fights with lions, (2) which individuals choose to participate, and (3) how the benefits of victory are distributed among cooperators and non-cooperators. We find that cooperative mobbing is more strongly influenced by lower costs (no male lions, more hyenas) than higher benefits (need for food). Individual participation is facilitated by social factors, both over the long term (close kin, social bond strength) and the short term (greeting interactions prior to cooperation). Finally, we find some direct benefits of participation; after cooperation, participants were more likely to feed at contested carcasses than non-participants. Overall, these results suggest that, when animals play dangerous cooperative games, selection favors flexible strategies that are sensitive to dynamic factors emerging over multiple time-scales.

## INTRODUCTION

Humans and other animals cooperate when the net benefits of cooperation exceed benefits accruing to individuals acting alone ^1^. One type of cooperation is collective action, where many individuals cooperate to gain group-level benefits ^2^. In animals, collective action includes both intra- and inter-specific conflicts, such as driving away predators or defending territory, offspring, or resources ^3^. The group-level cooperation that occurs during collective action is an emergent property of decisions made by individuals to cooperate or defect ^4,5^. Collective action problems arise when group members choose to pursue individual rather than group benefits; where defectors are able to enjoy the collective benefits of cooperators, “cheater” strategies can arise ^2^.

What drives an individual to cooperate, rather than defect, when faced with a collective action problem? Participation in collective action can yield important individual-level benefits, including: (1) acquisition or defense of resources ^1^; (2) kin-selected fitness benefits among highly related group members ^6^; and/or (3) other indirect benefits and social incentives, such as an enhanced reputation with potential coalition partners or mates ^7,8^. However, participation is usually costly, involving opportunity and energetic costs, and risk of injury or death ^9,10^.

Previous work has predominantly focused on collective action in homogenous animal groups ^11^, but individual and relational heterogeneity in social groups can strongly influence decisions regarding whether or not to participate in collective action ^12,13^. Theoretical modelling suggests that group members are more likely to participate when they can expect the biggest share of the benefits or rewards, can contribute for the lowest cost, or are the most capable (e.g., largest, strongest) ^14,15^. Other theoretical studies have demonstrated the importance of social network connections to successful collective action, especially in societies where social relationships are critical to fitness ^16,17^. However, empirical studies about collective action within heterogenous animal groups are still lacking.

Spotted hyenas (*Crocuta crocuta*) are an ideal study system in which to investigate collective action in heterogeneous social groups: they live in complex, differentiated societies, called clans, which are large (≤130 individuals), mixed-sex, fission-fusion groups ^18^. All clan members know one another individually, rear their cubs together at a communal den, and defend a common territory ^19^, but to avoid competition, clan-mates spend much of their time alone or in small subgroups ^20^. Due to female philopatry and male dispersal, most east African clans are composed of multiple matrilines of adult females, their offspring, and several adult immigrant males ^21^. Mean relatedness among clan-mates is very low ^22^. Each clan is structured by a strict linear dominance hierarchy, with natal animals outranking immigrants ^19^. Social rank has large fitness effects because it allows high-ranking group members to usurp food from clanmates ^23,24^.

Hyena clan-mates frequently cooperate during collective action in diverse contexts ^25^, including the collective mobbing of lions (*Panthera leo*) (Figure 1). Cooperative mobbing is a conspicuous example of collective action, which occurs when two or more individuals synchronously approach or attack a threat ^26^. Lions are spotted hyenas’ main competitors; these species use the same food resources and frequently kleptoparasitize one another ^27^. By mobbing lions, hyenas can overwhelm them and drive them away ^19^, increasing hyenas’ probability of feeding when competing with lions over food ^28^. Lions are larger and stronger than hyenas and represent a main source of mortality among hyenas ^27,29^. Mobbing lions is, therefore, very risky for hyenas, and – as it often results in benefits to both cooperating and defecting group members ^8,30^ – represents exactly the conditions under which cheating is expected to destabilize cooperation ^2,3^.

**Figure 1.**
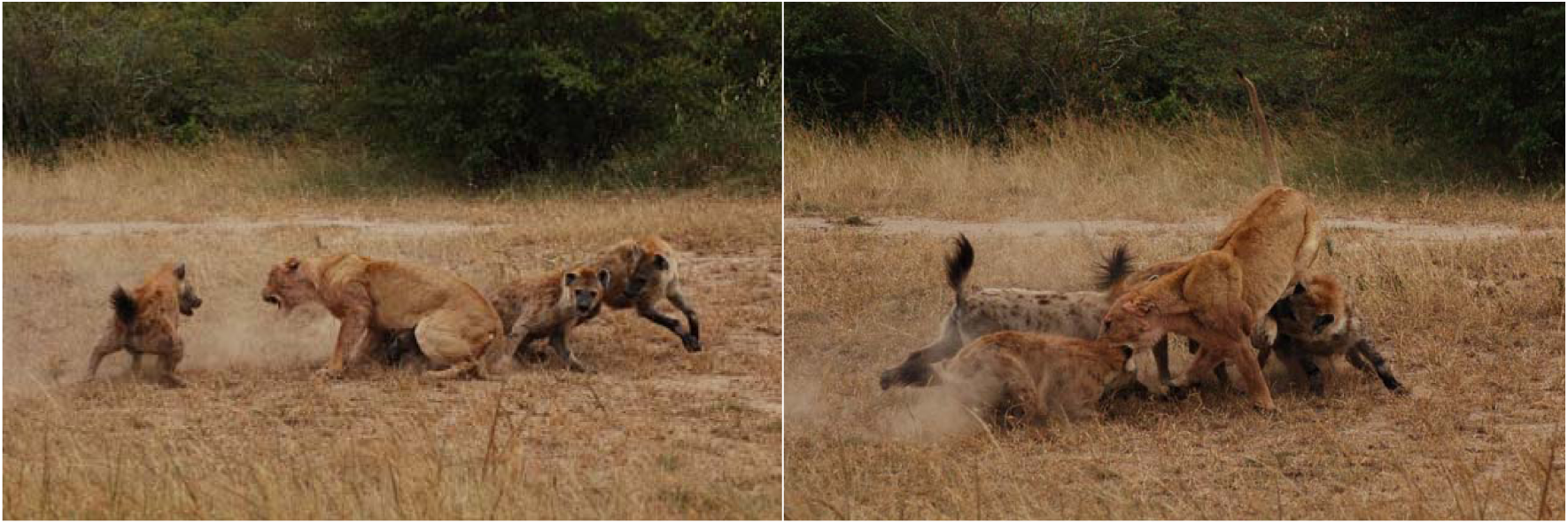
A group of four hyenas mobbing a lioness. Photos by Kate Yoshida.

Here, we aim to identify the mechanisms that drive cooperation in a complex society and to test theoretical predictions about collective action in heterogeneous groups. We focus on the collective mobbing of lions by wild spotted hyenas in Kenya and use a detailed, long-term dataset to investigate (1) when cooperative mobbing occurs, (2) who participates in cooperative mobbing, and (3) who benefits from it.

With respect to when mobbing occurs, based on past theoretical and empirical studies of inter-group conflict ^31^, we expected that both relative group size, and the ecological and social context in which lions and hyenas interact, would be critical variables. Specifically, we predicted that hyenas would be more likely to mob lions at valuable resources, such as the communal den or carcasses, especially when prey are scarce ^28,32^. We also predicted that hyenas would be more likely to mob lions when risks to individual hyenas are lower; namely, when male lions are absent and when the ratio of lions to hyenas is lower ^33,34^. Finally, we predicted that hyenas would be more likely to mob when they engage in affiliative interactions with groupmates or when they are more closely allied to the other individuals present ^35,36^.

With respect to who participates in mobbing, based on the theoretical modeling of Gavrilets and colleagues ^14,15^, we predicted that participants would be those with the lowest cost-benefit ratios. Hyenas would be more likely to participate, and less likely to defect, when they are high-ranking and thus have priority of access to any resources obtained via mobbing ^15,37^. We also predicted that hyenas would be more likely to participate when they are in top physical condition (i.e., prime-aged and good nutritional state), such that they can escape from lions more easily and thus bear a lower cost of participation ^14,38^. Based on theoretical studies showing the importance of social networks to successful cooperation ^16,17^, we also predicted that hyenas would be more likely to participate when their social allies or kin are present, as occurs in other socially complex species ^39,40^. Finally, hyenas engage in ritualized greeting behavior, which functions to promote cooperation and reinforce social bonds ^41^; we thus expected that occurrence of this affiliative behavior shortly before mobbing would increase an individual’s likelihood of participating ^36,42^.

With respect to who benefits from cooperative mobbing, we focused on potential individual-level food resource benefits of mobbing ^28^. We predicted that mobs would be more likely to occur when higher quality and/or larger food items are present ^43^. We also predicted that hungrier focal hyenas, as reflected by belly size, would be more likely to participate in mobbing when food is present ^44^. Most importantly, we predicted that hyenas who participate in cooperative mobbing would be more likely to obtain food ^28^.

## RESULTS AND DISCUSSION

To test these predictions, we built a series of logistic mixed effect models and performed model selection using AIC criteria on biologically relevant global models (Tables S1-S3) to determine which predictors were important determinants of mobbing occurrence, participation, and subsequent benefits.

### When does cooperative mobbing occur?

To identify the factors that influence whether a group of hyenas mob one or more lions, we used an occurrence dataset of all observation sessions (“sessions”) in which hyenas interacted with lions. Our logistic regression modeled whether or not a mob occurred during the session as a function of environmental and contextual factors that might affect mobbing occurrence (Model A in Table S1). This dataset contained 325 lion-hyena interaction sessions observed in 4 clans between 1988-2016.

Spotted hyenas mobbed in 41.8% (n = 136) of these sessions, with a median of 2 mobs per session (mean 3.1, range 1-40) and a median of 4 hyenas per mob (mean 5.1, range 2-16). Hyenas mobbed during 44% of interaction sessions at carcasses, at 35% of interaction sessions at active dens, and at 39% of interaction sessions away from either of these resources.

In our model of mobbing occurrence (Model A: n = 321 sessions; Figure 2; Table S4), mobbing was more likely to occur when more hyenas were present (β-hyenas = 0.87, p < 0.001) and when male lions were absent (β-male lions = −0.73, p = 0.014). Counter to our expectations, local prey density was positively correlated with the probability of mobbing (β-prey = 0.27, p = 0.038). Increasing numbers of individuals who engaged in greeting behavior (“greeters”) during the session also increased the predicted probability of mobbing (β-greeters = 0.53, p = 0.017). However, a near-significant interaction between number of hyenas present and number of greeters (β-hyenas x greeters = −0.32, p = 0.061) indicated that greetings may facilitate mobbing behavior when only a few hyenas are present, but do not appear to affect mobbing behavior when many hyenas are present. A near-significant interaction between male lion presence and number of greeters (β-male lions x greeters = 0.67, p = 0.059) indicated that greetings have a larger positive effect on mobbing occurrence when male lions are present than when they are absent. Session length, session context, number of lions present, and mean association index of hyenas present were not included in the top model or any model within 6 AIC of the top model.

**Figure 2.**
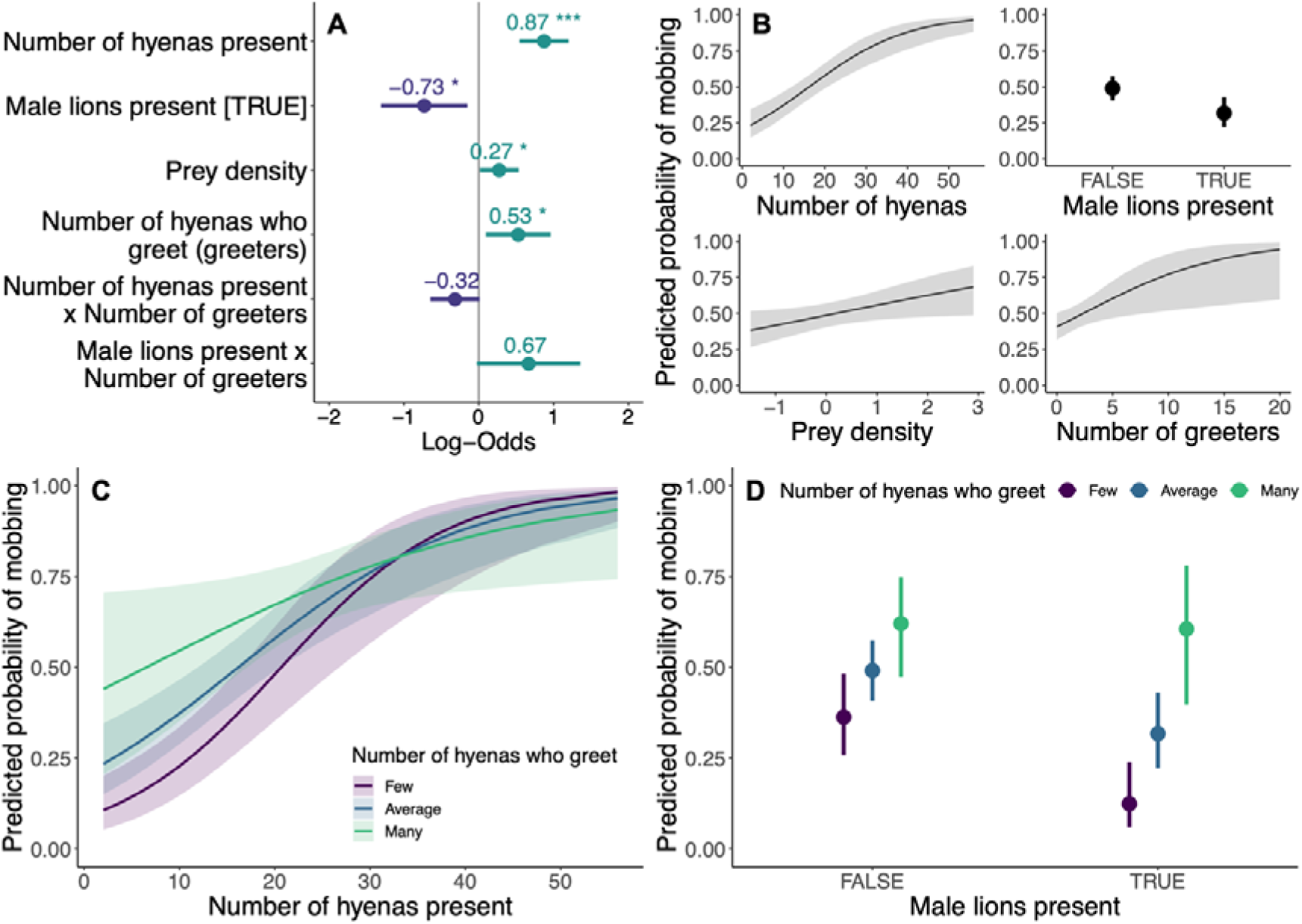
Top model of the predicted probability of mobbing occurrence in sessions where lions and hyenas interact (Model A: n-sessions = 321). **A.** Dots depict coefficient estimates, lines depict 95% confidence intervals, and asterisks depict significance at the following p-values: * = 0.05; ** = 0.01; *** = 0.001. **B-D.** Lines (or dots) depict estimated marginal means and shaded areas (or vertical lines) depict 95% confidence intervals.

Overall, our results demonstrate that the decision to mob lions is more strongly influenced by the situational risks of mobbing than the potential benefits: hyenas were most likely to mob in sessions where risk was reduced by more hyenas being present and male lions being absent, regardless of prey abundance or whether there were resources present to defend. Our results indicate that hyenas attend only to the presence or absence of male lions as a source of risk, as they did not otherwise alter their mobbing behavior based on the number of lions present (Models B and C in Tables S1 and S4), suggesting that the ratio of lions to hyenas may be less important than previously thought ^33,34^. Finally, we found that greetings were associated with increased mobbing occurrence, particularly when the situational risks were higher (i.e., fewer hyenas or male lions were present). This accords with prior studies suggesting that greetings promote cooperation and reinforce social bonds^41^. Our results imply an additional critical function for greetings as a coordination mechanism allowing hyenas to achieve collective action.

### Who participates in cooperative mobbing?

To understand cooperative mobbing at the individual level, we examined the factors predicting an individual’s participation in mobbing, given that a mobbing event occurs. To do this, we restricted our dataset to lion-hyena interaction sessions where mobbing occurred and where the identities of more than 90% of mobbing participants were known. We then built a series of logistic mixed-effect models where we modeled whether a focal hyena chose to cooperate (mob) or defect (not mob) during a mobbing event where they were present. We included key demographic and social factors with the potential to influence mobbing participation as fixed effect covariates for each focal hyena in these models (Models D-G in Table S2). All models included random intercept covariates of hyena identity and of mobbing event nested within session.

This participation dataset consisted of 4740 mob-hyena combinations, with 492 unique hyenas present for 344 total mobs during 119 observation sessions involving lions and hyenas. In 33% (n = 1577) of mobbing opportunities, focal hyenas participated in mobs, while in the remaining 67% (n = 3163) of mobbing opportunities, focal hyenas were present, but defected. Of the 492 unique hyenas, 44 individuals always mobbed (in a range of 1-5 mobs), and 189 individuals always defected (in a range of 1-44 mobs). The remaining 259 hyenas mobbed in a median of 33% (mean = 38%; range = 2-94%) of mobbing opportunities (median = 9; mean = 14.8; range = 2-94 mobs).

In our overall participation model (Model D: n = 4383 mob-hyena combinations; Figure 3; Table S4), females were more likely to mob than males (β-male = −1.04, p < 0.001). Focal individuals of age 7.6 years (range 0.2-21.2 years) were most likely to mob (β-age = 0.72, p < 0.001; β-age^2^ = −0.40, p < 0.001). Because of these clear differences based on age and sex, we divided all subsequent analyses by sex and age class. Spotted hyenas reach reproductive maturity at 2 years of age ^45^, so individuals were either juveniles (< 2 years; see Supplementary Material) or adults (>= 2 years).

**Figure 3.**
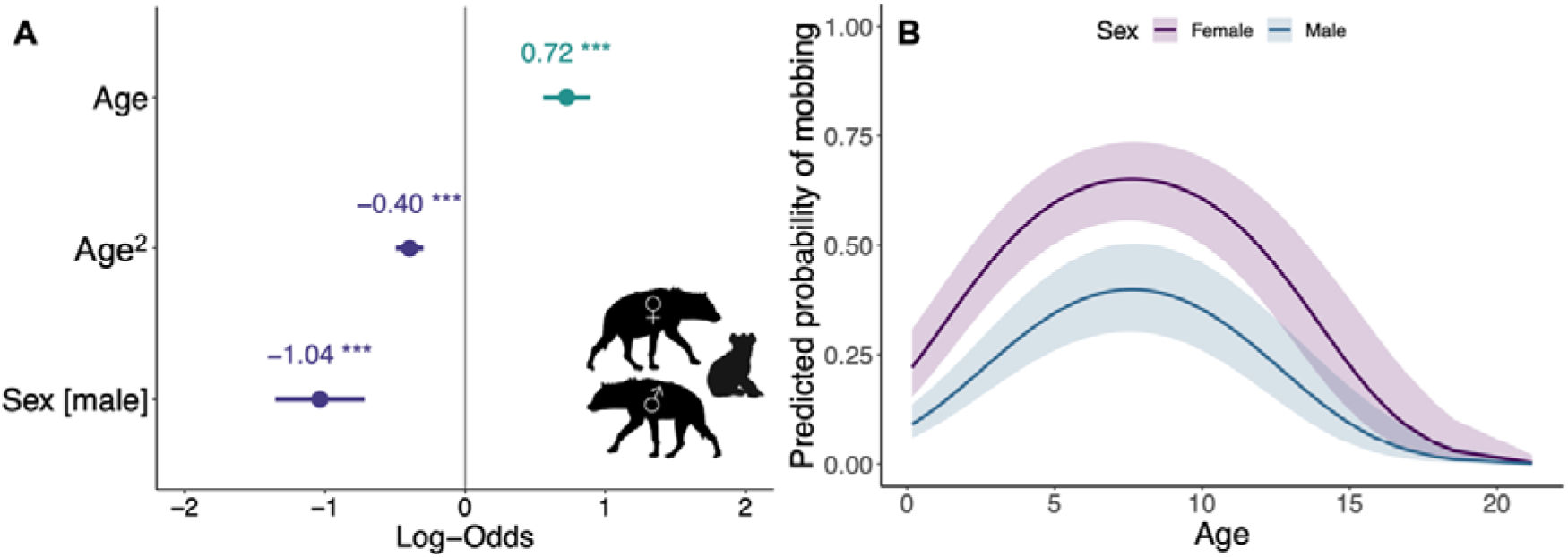
Top model for probability of mobbing participation by all hyenas (Model D: n-focal hyenas = 4383; n-sessions = 117; n-mobs = 342; n-unique hyenas = 431). **A.** Dots depict coefficient estimates, lines depict 95% confidence intervals, and asterisks depict significance at the following p-values: * = 0.05; ** = 0.01; *** = 0.001. **B.** Lines depict estimated marginal means and shaded areas depict 95% confidence intervals.

In our adult female participation model (Model E: n = 2280 mob-hyena combinations; Figure 4; Table S4), focal females that were 6.7 years old (range 2.0-21.2 years) were most likely to mob (β-age = 0.08, p = 0.410; β-age^2^ = −0.13, p = 0.014). Social rank was included in the top model but was not significantly associated with mobbing behavior (β-rank = 0.17, p = 0.113). Here, as in the model of mobbing occurrence, greetings strongly promoted mobbing behavior; females that engaged in greeting behavior during the 5 minutes before the mobbing event occurred were more likely to mob than those that did not greet (β-greeted = 1.17, p < 0.001). A significant interaction between greeting behavior and social rank revealed that greeting more strongly promoted mobbing for low-than high-ranked females (β-greeted x rank = −0.75, p = 0.009). Focal females were more likely to mob if other participants were their more frequent associates (β-association index = 0.39, p = 0.004). Again, there was an interaction between frequency of association and social rank: association strength with participants was correlated with higher mobbing probability for high-but not low-ranking individuals (β-association index x rank = 0.21, p = 0.024). Lastly, focal females were more likely to mob if they were related to a larger proportion of the current participants (β-maternal relatedness = 0.23, p = 0.013). Reproductive state was not included in the top model or any model within 6 AIC of the top model.

**Figure 4.**
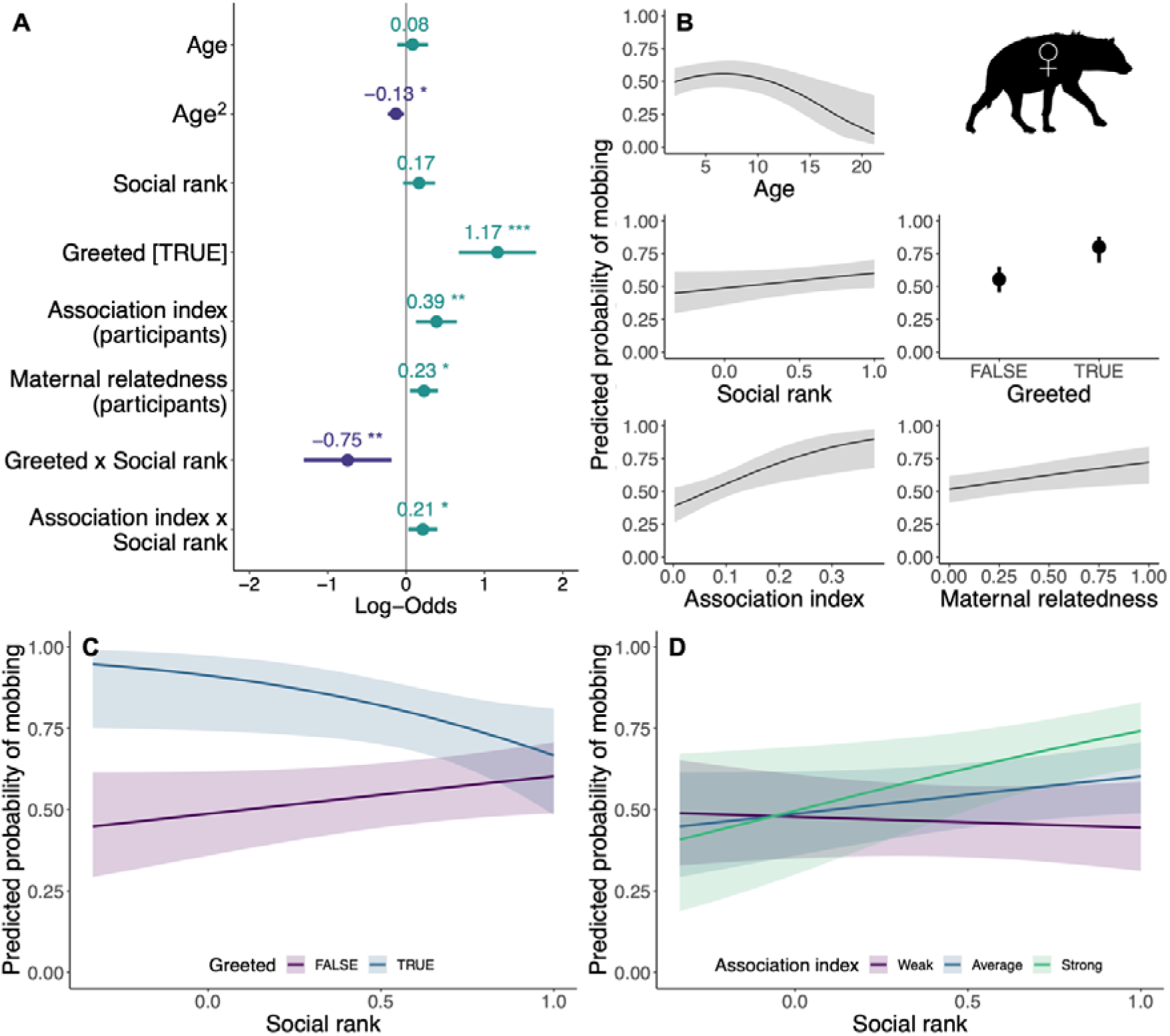
Top model of the predicted probability of mobbing participation by adult female focal hyenas (Model E: n-focal hyenas = 2280; n-sessions = 109; n-mobs = 323; n-unique hyenas = 169). **A.** Dots depict coefficient estimates, lines depict 95% confidence intervals, and asterisks depict significance at the following p-values: * = 0.05; ** = 0.01; *** = 0.001. **B-D.** Lines (or dots) depict estimated marginal means and shaded areas (or vertical lines) depict 95% confidence intervals. **D.** Association index with participants was analyzed as a continuous variable but is depicted categorically for illustrative purposes.

In our adult male participation model (Model F: n = 893 mob-hyena combinations; Figure 5; Table S4), focal males that were 6.2 years old (range 2.0-16.9 years) were most likely to mob (β-age = 0.12, p = 0.616; β-age^2^ = −0.37, p = 0.025). Higher-ranking males were more likely to mob than their lower-ranking counterparts (β-rank = 0.97, p < 0.001). Focal males that were close associates of the current participants were more likely to participate in that mob than males that were weakly associated (β-association index = 0.36, p = 0.045). Interestingly, in contrast to models of mobbing occurrence and of female participation in mobbing, male mobbing was not facilitated by greeting behavior. Neither dispersal status nor whether the focal hyena greeted during the 5 minutes before the mob formed were included in the top model or any model within 6 AIC of the top model.

**Figure 5.**
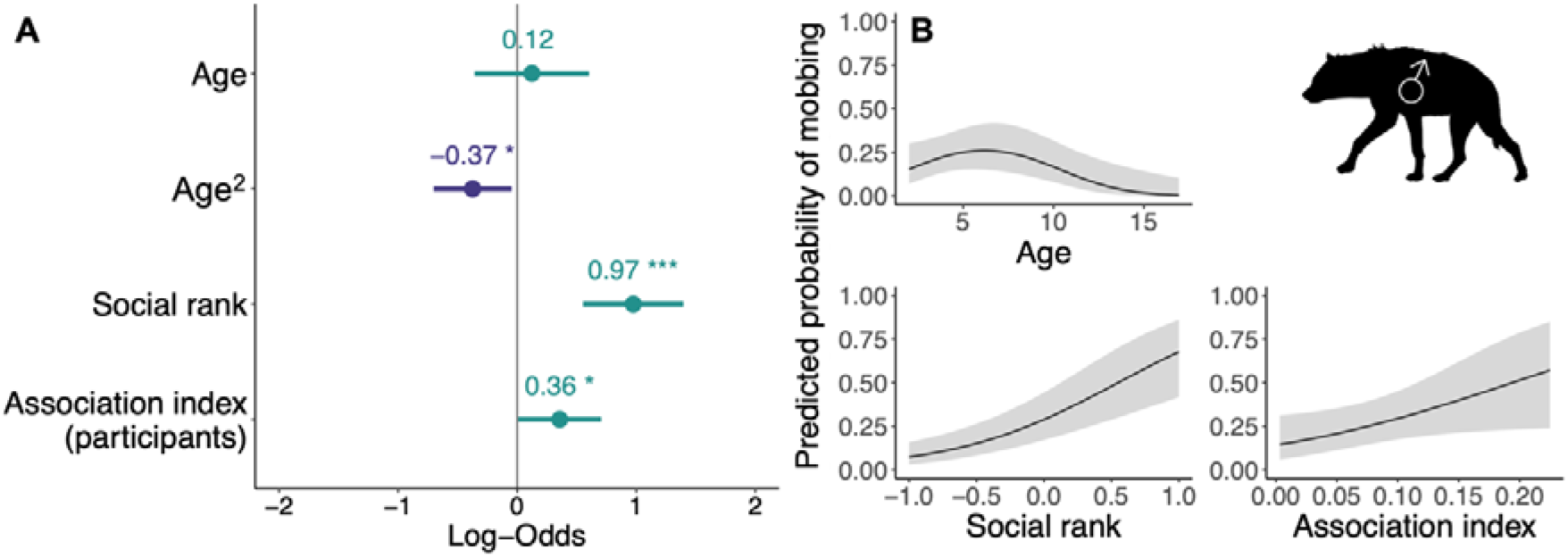
Top model of the predicted probability of mobbing participation by adult male focal hyenas (Model F: n-focal hyenas = 893; n-sessions = 101; n-mobs = 288; n-unique hyenas = 124). **A.** Dots depict coefficient estimates, lines depict 95% confidence intervals, and asterisks depict significance at the following p-values: * = 0.05; ** = 0.01; *** = 0.001. **B.** Lines depict estimated marginal means and shaded areas depict 95% confidence intervals.

Our participation models revealed that characteristics suggesting a stronger individual, such as being female (the larger sex), prime-aged, and higher-ranking, predicted a higher likelihood of mobbing. Mobbing decisions by females were sensitive to greeting behavior and to longer-term social factors such as associative relationships, social rank, and kinship. Adult males also participated in mobbing, although less frequently than adult females. Despite having weaker social bonds within the group ^46,47^, adult male mobbing behavior was also correlated with variation in social relationships, including dominance rank and association strength. Overall, our results suggest that hyenas’ decisions to cooperate in mobbing are strongly affected by the local social environment, both short-term interactions (greeting) and long-term relationships (association, rank, kinship).

### Who benefits from cooperative mobbing?

Past research suggested that mobbing behavior might facilitate the acquisition or defense of food resources. To investigate this, we restricted our dataset to observation sessions with food present (Table S5), and further restricted our participants to focal adult hyenas, as juvenile resource acquisition and defense are strongly dependent on adult support ^48,49^. We then built a series of logistic models where we modeled the probability of mobbing occurrence, mobbing participation, and benefits to participants as a function of food-related variables such as carcass size and freshness, individual nutritional state, and individual feeding after mobbing events (Models H-K in Table S3).

We first inquired whether hyenas are more likely to mob to obtain or defend larger or fresher food resources. Our top model of the occurrence of mobbing at sessions with food (Model H: n = 218 sessions; Table S4) did not include the term for carcass size but did include the term for carcass freshness (β-freshness = −0.32, p = 0.443), although the effect was non-significant. This suggests that mobs are equally likely to occur across all food sessions, regardless of carcass size or quality.

We next tested whether hyena nutritional state, indicated by individual belly size at the start of the session, affected mobbing participation at carcasses of different sizes. In the model of adult hyena mobbing participation during sessions with food (Model I: n = 407 mob-hyena combinations; Figure ED1; Table S4), “obese” individuals were marginally less likely to mob than either “fat” or “normal” individuals (Tukey post-hoc test for belly size: [obese - normal]: β = −2.50, p = 0.060; [obese - fat]: β = −2.54, p = 0.057), although there was no difference in mobbing participation between “normal” and “fat” individuals ([fat - normal]: β = 0.04, p = 0.990). Focal individuals were also less likely to mob at the largest carcasses (Tukey post-hoc test for carcass size: [extra-large - medium]: β = −3.75, p = 0.027; [extra-large - large]: β = −2.63, p = 0.048; [large - medium]: β = −1.12, p = 0.493). Hyenas’ age (β-age = 0.45, p = 0.007; β-age2 = −0.19, p = 0.033) and social rank (β-rank = 0.73, p < 0.001) also significantly affected their probability of mobbing, as shown in earlier models.

Finally, we tested whether participants were more likely than non-participants to obtain food after mobbing events. In our model of the probability of adult hyenas feeding in the 5 minutes after a mob occurred (Model J: n = 1049 mob-hyena combinations; Figure 6; Table S4), focal individuals that mobbed were significantly more likely to feed than individuals that defected (β-participant = 0.56, p = 0.006), even after controlling for age (β-age = 0.08, p = 0.668; β-age2 = −0.26, p = 0.005) and social rank (β-rank = 0.42, p = 0.006). However, the model of hyenas feeding at any point during the session (Model K: n = 673 session-hyena combinations; Table S4) did not include the term for whether or not a focal hyena mobbed during the session, nor did any models within 6 AIC of the top model.

**Figure 6.**
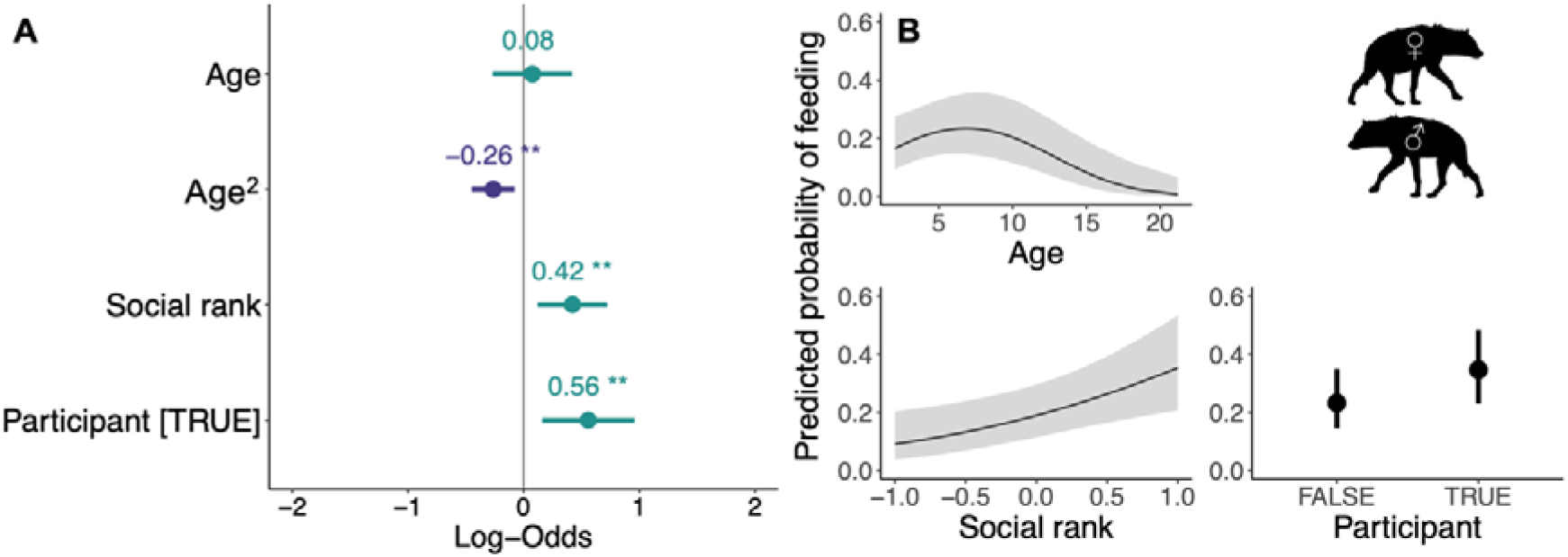
Top model of the predicted probability of the focal hyena feeding during the five minutes immediately after a mob (Model J: n-focal hyenas = 1049; n-sessions = 41; n-unique hyenas = 185). **A.** Dots depict coefficient estimates, lines depict 95% confidence intervals, and asterisks depict significance at the following p-values: * = 0.05; ** = 0.01; *** = 0.001. **B.** Lines (or dots) depict estimated marginal means and shaded areas (or vertical lines) depict 95% confidence intervals.

Our results indicate that mobbing increases access to food for spotted hyenas, but that hyenas generally do not adjust their mobbing behavior based on potential food rewards. One exception to this pattern was that hyenas were less likely to mob at extra-large carcasses such as hippos and elephants; these carcasses last for days in our study area ^50^, so it may be unnecessary to risk mobbing when simply waiting will yield rewards ^43^. Additionally, obese hyenas were less likely to participate in mobbing than thinner individuals, perhaps because they are already satiated or because their obesity may impair their movement (Figure ED2). Our past research suggested that mobbing increases the probability of any one hyena in the session getting to feed ^28^. Here we extend this finding by showing that hyenas who mob during contests with lions over food were more likely to feed in the 5 minutes after the mob. Although this benefit was short-lived, food obtained by participants during or immediately after mobbing could be substantial, as hyenas can consume enormous quantities of meat extremely quickly ^19^.

## CONCLUSION

### Theoretical predictions

Here, our overarching goal was to deepen our understanding of collective action in complex societies ^51^ and to test theoretical predictions about collective action in heterogeneous groups. Overall, our results support theoretical work suggesting that participants in collective action are often those group members with the lowest cost-benefit ratios ^14,15^. Interestingly, we also found that participation in mobbing was more sensitive to the potential costs of participation than the potential benefits of success. Furthermore, our finding that long-term social ties were associated with mobbing supports theoretical work demonstrating the importance of social network connections to successful group cooperation ^16,17^. Importantly, our work adds a third critical component to successful collective action: short-term prosocial behaviors. Greetings promoted both mob occurrence and participation across age classes. These affiliative behaviors could be an important precursor to risky cooperative behavior via a “raising the stakes” model of cooperative partnership formation ^52^.

### Why act collectively?

Using a dramatic example of risky cooperative mobbing against a dangerous predator and competitor, we demonstrate how the coordination of collective action is contextualized within the broader environment of a society characterized by many different types of social relationships. The striking variation among individuals and relationships in such groups complicates decision-making regarding whether or not to cooperate. However, across contexts, we found that short-term prosocial behaviors boosted individual and group-level cooperative tendencies, sometimes allowing groups to achieve collective action that would otherwise not occur. The benefits of engaging in this collective action were harder to pin down. Although we found some evidence that individuals gained direct benefits from mobbing lions, we found only mixed support for the idea that hyenas adjust their mobbing behavior in response to these potential benefits, and a quarter of mobbing events occurred in the absence of any obvious immediate reward. Overall, we found that, when playing cooperative games, hyenas, and perhaps also most other animals living in complex societies, choose cooperative strategies flexibly. Instead of “cooperators” and “defectors,” complex societies in nature are composed of individuals exploiting the flexibility of behavior to cooperate or defect depending on dynamic factors that emerge over multiple time-scales. This suggests that social selection may favor individuals that attend to the social characteristics of their group-mates so they can safely navigate risky collective action together.

## METHODS

From 1988-2016, we monitored four clans of wild spotted hyenas in the Maasai Mara National Reserve, Kenya. We monitored one clan from 1988-2016 and three clans from 2008-2016. We monitored clans daily during two observation periods, from 0530-1030 and from 1600-2100. When we encountered a subgroup of one or more hyenas, we initiated an observation session (“session”) and recorded the identities of all hyenas present within a 200 m radius, using their unique spot patterns and ear damage to recognize individuals. We also recorded the number, sex, and age class of all lions found ^53^. Sessions lasted from 5 minutes to several hours and ended when behavioral interactions ceased, and observers left that individual or group. Using all-occurrence sampling ^54^, we recorded arrivals and departures of individual hyenas, agonistic interactions, and greetings. Greetings are affiliative interactions occurring when two partners stand parallel to one another but facing in opposite directions to sniff the other’s anogenital region ^19^. We also performed scan-sampling ^54^ every 20 minutes throughout each session to document change in hyenas present.

In our population, lions and hyenas co-occurred in an average of 4 sessions per clan per month, and the two species interacted (directed behavior at one another) in 44% of those co-occurrence sessions ^28,55^. Throughout each session involving both lions and hyenas, we recorded all mobbing events using all-occurrence sampling. We operationally defined “mobbing” as a group of two or more hyenas, usually side-by-side and within 1 m of one another, with tails bristled over their backs, approaching within 10 m of at least one lion (Figure 1; Lehmann et al., 2017). In association with each mobbing event, we recorded the identities of all participating hyenas and the number, sex, and age class of the lions being approached.

Throughout each lion-hyena session in which a kill or carcass was present, we recorded hyena feeding behavior. Because lion-hyena sessions are often very chaotic (and thus the ability of the observer to record feeding behavior varies), we created a simple feeding dataset of one-zero sampling for each hyena present at each session. For each minute of each session, we recorded whether or not a focal hyena was observed feeding. To be conservative, any minutes in which a focal hyena was not directly observed feeding were coded as “not feeding.”

Because hyena societies are fission-fusion and most individuals spend the majority of their time alone or in small subgroups, we measured the strength of social relationships among individuals by calculating association indices ^56^. Association indices were calculated for each dyad in each session using R package *asnipe* ^57^ based on patterns of association over the previous 365 days. Thus, we calculated the association index within the dyad of hyenas A and B as (A+B_together_) / [(A_without B_) + (B_without A_) + (A+B_together_)] where (A+B_together_) represents the number of sessions in which A and B were both present, (A_without B_) represents the number of sessions in which A was observed but B was not present, and (B_without A_) represents the number of sessions in which B was observed but A was not present.

### Model predictors

#### Variables calculated for each observation session

*Session length:* The length of the session in minutes.

*Session context:* We assigned each session to one of three contexts: “food” sessions occurred within 200 m of a kill or carcass, “den” sessions occurred within 200 m of an active hyena den, and locations of all remaining sessions were categorized as “other” sessions ^58^, in which hyenas were usually resting or traveling.

*Prey density:* For each session, we calculated the current prey density on a monthly basis. We monitored prey availability during biweekly surveys by counting all wild herbivores within 100 m of 2-3 line transects (1.5-5.4 km long) in each clan territory ^28,59^. Prey density was calculated as prey counted per square kilometer based on the number of animals sighted during line transect surveys. For each month of our study, we calculated the prey density within the territories of each of our study clans and used the monthly number of standard deviations above or below the yearly mean prey density to determine standardized prey availability for each clan during each month of study ^28^.

*Number of hyenas present:* The total number of hyenas present in the session.

*Number of lions present:* The total number of lions present in the session.

*Male lions present (Y/N):* Whether or not adult male lions were present in the session.

*Number of male lions present:* The total number of adult male lions present in the session.

*Number of hyenas who greet (greeters):* For each session, we quantified the amount of affiliative behavior observed as the total number of individuals present who engaged in greeting behavior during that session.

*Mean association index:* For each session, we quantified the strength of social ties between individuals present in that session as the mean association index of all dyads present in that session.

*Carcass freshness:* At food sessions, we categorized the carcass as “fresh” (prey was recently killed) or “old” (prey was killed over 24 h earlier). Past work referred to fresh carcasses as “kills” and old carcasses as “carcasses” ^28^.

*Carcass size:* At food sessions, we recorded the species, sex, and age class (where possible) of the carcass when first seen. Based on observer descriptions, carcass size was later categorized by prey species age and weight ^60^ as small (< 20 kg), medium (20-100 kg), large (100-500 kg), and extra-large (> 500 kg; Table S6). Small and medium categories were combined due to the limited sample size for small carcasses.

#### Variables calculated for each focal hyena at each mobbing event

*Age:* We estimated cub birthdates (to ± 7 days) based on their appearance when first seen ^61^. We considered individuals of both sexes to be juveniles until 24 months of age, after which they were considered adults ^45^. All immigrant males were classified as adults; whenever possible, their birthdates (to ± 6 months) were estimated via tooth measurements obtained during routine immobilizations or necropsies ^62^. Here we calculated the age in years of each focal hyena on the date of each session, and we fit a second-order polynomial for age for all adults because very young and very old hyenas may be less likely to mob.

*Sex:* Hyenas were sexed based on the morphology of the erect phallus ^63^.

*Social rank:* The social rank of each hyena was determined based on the occurrence of submissive behavior during dyadic agonistic interactions ^64^. For each clan, separate social ranks were calculated annually for adult females and adult males using the MatReorder method in R package *DynaRank* ^65^. For each year, the sex-specific hierarchies were then combined, with all adult females dominant to all adult immigrant males ^19^, and an annual standardized rank was calculated for each adult within each clan. Juveniles in our dataset were assigned the same rank as their mother for each year until they either became adults at age 2 (females) or dispersed between ages 2-5 (males). Males who did not disperse were added to the top of the adult immigrant male hierarchy when they reached 5 years of age ^66^. Here we calculated the social rank of each focal hyena as its social rank in the calendar year of the session.

*Reproductive state (females):* Reproductive states of all adult females were continuously monitored to determine periods of pregnancy and lactation ^61^. A female was considered “nulliparous” until her first parturition, determined by direct observation of her first litter or by the observation of pink scar tissue on the posterior surface of her phallus, which tears during parturition ^67^. Conception dates were determined by subtracting a 110-day gestation period from a cub’s date of birth ^19^ and/or by observations of fresh tears in a female’s phallus, indicating recent parturition. A female was considered “pregnant” from conception until a cub’s date of birth, and “lactating” from the day after that birth until the latest weaning date of a cub from that litter. Weaning dates for each cub were calculated based on observations of nursing conflicts and observations of cubs subsequently seen with their mother when no nursing occurred ^61^. Cubs that did not successfully wean were considered “dead before weaning,” and their disappearance date was substituted for their weaning date in the calculation of maternal reproductive state. A focal female that was neither nulliparous, pregnant, nor lactating on the session date was assigned to reproductive state “other.”

*Dispersal status (males):* Dispersal status was considered as either “immigrant” if a male had immigrated from his natal clan into the study clan, or “natal” if the male was born in the study clan, regardless of age. Dispersal status is a good proxy for reproductive status in males because immigrant males sire 97% of all juveniles born in the clan ^68^, and immigrant males have higher testosterone and higher ejaculate quality than do age-matched adult natal males ^69,70^.

*Greeted (Y/N):* We measured individual affiliative behavior as whether or not the focal hyena had engaged in greeting behavior with a group-mate during the five minutes before the mob. Five minutes is frequently used as a window to measure the effects of behavior in this species ^71^.

*Association index with participants:* We measured the strength of social ties between a focal hyena and the mobbing individuals to evaluate the effect of relationship strength on the probability that a focal hyena would mob. For each focal hyena, we calculated the mean association index between the focal individual and all mobbing individuals; in other words, we averaged the association index of all dyads involving the focal individual and each mobbing individual.

*Maternal relatedness with participants:* Throughout our study, we established maternity for all resident hyenas based on nursing associations ^61^. To evaluate the effect of kinship on the probability that an individual would mob, we calculated the proportion of participants to whom the focal individual was related. Because many sires are immigrant male hyenas for whom we currently lack paternity data, relatedness was evaluated solely on the basis of maternal kinship. For each focal hyena, we calculated the proportion of participants to whom the focal individual was closely maternally related as the number of participants to whom that individual was either a mother, offspring, or sibling, divided by the total number of participants.

*Belly size:* We recorded the belly size of all focal adult hyenas upon first sighting in a session as one of four states: “gaunt” hyenas were very skinny with hipbones protruding; “normal” hyenas were fit but not fat; “fat” hyenas had a big full belly; and “obese” hyenas had a truly monstrously giant belly ^72^. Gaunt and normal categories were later combined due to the tiny sample size of gaunt hyenas.

### When does cooperative mobbing occur?

Here, we restricted our dataset to observation sessions in our four study clans where lions and hyenas interacted. We operationally defined interspecific interactions as occurring when lions and hyenas directed behavior at one another and/or when lions and hyenas approached within 10 m of one another ^28^. We further filtered to sessions with field notes of high-enough quality to be certain that all mobbing events were recorded if they were observed. Finally, we filtered to sessions where at least 2 hyenas were present because, by definition, multiple hyenas are required for a mob to occur. We fit a logistic regression where our response variable was binomial: whether or not a mob occurred during that session. Fixed effect covariates included key environmental and contextual variables with the potential to affect mobbing occurrence (Model A in Table S1). We included interactions between session length and the number of hyenas present, and between session length and the number of hyenas that greet (greeters), to control for the possible correlation between observation time and number of hyenas or greetings observed. We included interactions between number of hyenas present and total number of lions present, and between number of hyenas present and male lions present based on past work indicating that the ratio of lions to hyenas can affect mobbing behavior ^33,34^. We included interactions between hyena and lion variables (number of hyenas present, number of lions present, male lions present) and social variables (number of greeters, mean association index) to test whether social behavior could help overcome the barriers to mobbing we documented earlier ^28^. No random effects were included in this model; clan was considered as a random intercept but was dropped as it explained no variance.

### Who participates in cooperative mobbing?

Here, we restricted our dataset to observation sessions where mobbing occurred and where the identities of more than 90% of mobbing participants were known. For each mob during these sessions, we determined which hyenas were present when the mob occurred based on the arrival and departure times of all hyenas in the session. Each focal hyena present during a mobbing event was coded as either a participant (“participant”) or non-participant (“defector”) for that particular mobbing event. We then assigned relevant demographic, physiological, and social variables to each focal hyena: we assigned an age, social rank, reproductive state (females), and dispersal status (males) to each focal hyena present. We also assigned social context measures to each focal hyena present, including whether or not the focal hyena had greeted in the five minutes prior to a mob (“greeted”), the average association index between the focal hyena and other participants (“association index”), and the proportion of participants to which the focal hyena was closely related (i.e., mother, sibling, or offspring of the focal hyena; “maternal relatedness”).

To investigate hyena participation in cooperative mobbing events, we fit a series of logistic mixed-effect models where our response variable was binomial: whether or not the focal hyena participated in that mob. Fixed effect covariates included key demographic and social variables with the potential to affect mobbing participation (Models D-G in Table S2). All models included random intercept covariates of hyena identity and of mob nested within session. Clan was not included as a random intercept because it explained only 2.2% of the variance in participation (intraclass correlation coefficient = 0.022).

We built a series of logistic mixed-effect models to test the effects of different variable sets on specific categories of hyenas.

*Preliminary analysis of all hyenas.* The first model (Model D in Table S2) included all hyenas and included age and sex to identify broad differences between age and sex classes.

*Female participation model.* The second model (Model E in Table S2) was restricted to all adult females (age > 2 years) and included key demographic and social factors with the potential to affect mobbing participation in adult females. We included interactions between social rank and other variables because social rank critically structures hyena social relationships ^73^.

*Male participation model.* The third model (Model F in Table S2) was restricted to all adult males (age > 2 years) and likewise included key demographic and social characteristics with the potential to affect mobbing participation in adult males. We included interactions between social rank and other variables. We were not able to include the term for maternal relatedness in this model because many of these individuals were immigrant males for which we do not currently have relatedness data. We were also unable to include an interaction between age and social rank due to its collinearity with social rank.

*Juvenile participation model.* Our fourth model (Model G in Table S2) was restricted to all juveniles (age < 2 years) and included key demographic and social factors with the potential to affect mobbing participation by juvenile hyenas. We also included three interaction terms, age by sex, age by social rank, and sex by social rank.

To ensure that we were measuring the effect of affiliative social interactions and not just that of social interactions more generally, we re-ran top models that included a term for whether or not a hyena greeted to also include a term for whether or not an individual engaged in an aggressive interaction in the five minutes prior to the mob occurring. In none of these models was the aggression term included in the top model, whereas the affiliative term remained in top models, confirming that our greeting measure captures the effect of affiliation specifically and not of social interactions more generally.

### Who benefits from cooperative mobbing?

To investigate potential resource benefits of mobbing, we fit four logistic mixed-effect models (Models H-K in Table S3). For all analyses of resource benefits, we restricted our dataset to observation sessions with food present, and further restricted our participants to focal adult hyenas (age > 2 years), as juvenile resource acquisition and defense are strongly dependent on adult support ^48,49^. If hyenas mob to obtain or defend food resources, we predicted that mobs would be more likely to occur at sessions where higher quality (fresher) and/or larger food items were present (Model H in Table S3). Here, we modified our global model of the probability of mobbing occurrence (Model A in Table S1) by including terms for food quality (“carcass freshness”) and size (“carcass size”).

In our second model (Model I in Table S3), we predicted that hyenas that were hungrier, or those in a poorer nutritional state, would be more likely to participate in mobbing at sessions with food. Here, we fit a logistic mixed-effects model with a binomial response variable: whether or not the focal hyena mobbed during the session. We restricted our analysis to focal adult hyenas during sessions in which observers had recorded at least one non-normal belly size to create more even categorical distributions for belly size. This model included the following fixed effects: age, sex, social rank, belly size, carcass size, and carcass freshness. We also included interactions between social rank and belly size and between social rank and carcass size because of the large effect that social rank has on resource acquisition ^23^.

Lastly, we predicted that hyenas who participate in mobbing would be more likely to obtain food, both immediately after the mob and during the session overall. For these analyses, we restricted our dataset to mobs (Model J in Table S3) or sessions (Model K in Table S3) where at least one hyena fed, and we coded each hyena present as either a mobbing participant or defector. We built two logistic mixed-effects models to test these predictions, where the response variable was binomial: whether or not that hyena fed. Both models included the following fixed effects: focal hyena age, sex, social rank, and participant status, carcass freshness and size, and interactions between social rank and participant and between participant and carcass size. Model J investigated the probability of the hyena getting food within 5 minutes after the mob and included a fixed effect of whether or not the focal hyena participated in that mob. Here, for each mob, our response variable was whether or not the focal hyena fed in the five minutes following the mob. We removed mob identity as a random effect from the global model (Model J) because it explained no variance. Model K investigated the probability of a hyena getting food during the session overall and included a fixed effect of whether or not the focal hyena mobbed during the session. Here, for each session, our response variable was whether or not the focal hyena fed anytime between the first mobbing event and 30 minutes after the final mobbing event. We excluded later feeding data to reduce feeding observations due to hyena turnover at the carcass as some hyenas become satiated, and we used 30 minutes as our cut-off because a group of hyenas can reduce a large carcass to bones in under 30 minutes ^19^. We also removed hyena identity as a random effect from the global model (Model K) because it explained no variance.

### Statistical analysis

All analyses were conducted using R Version 4.1.2 and R Studio Version 2021.09.0. We first performed data exploration by investigating outliers, distributions and collinearity ^74^. We tested all global model predictors for multicollinearity using both correlation coefficients and variance inflation factors (VIFs), and we removed collinear predictors until none were collinear, with all correlation coefficients ≤ 0.7 and all VIFs ≤ 3 ^75^. All numeric model predictors were z-score standardized immediately before modeling using the scale function in R to allow comparison of coefficients ^75^. We used R package *glmmTMB* ^76^ to build all models, and we performed model selection on the global model using the dredge function in R package *MuMIn* ^77^. The top models, as determined by AIC criteria, are depicted in the figures and tables here and in our supplementary information. All top models were visually inspected to confirm assumptions of multicollinearity, normality of residuals, normality of random effects, heteroscedasticity, and homogeneity of variance using R package *performance* ^78^ and R package *DHARMa* ^79^. We also used R package *DHARMa* to inspect all groups and observations for disproportionate influence in our models, but none warranted exclusion. Between-group comparisons were conducted using Tukey post-hoc tests for multiple comparisons of means in R package *multcomp* ^80^. Forest plots were created using R package *sjPlot* ^81^ and all other plots were created using the ggpredict function in R package *ggeffects* ^82^ to obtain predicted values and R package *ggplot2* ^83^ to create the plots from those values.

## DATA & CODE AVAILABILITY

The data and source code are available through a GitHub repository (https://github.com/tracymont/hyena_mobbing), which will be made public upon publication.

## ACKNOWLEDGEMENTS

We first and foremost thank all past and present members of the Mara Hyena Project for their work collecting and curating the data presented here. We thank Meghan Bugaj, Cameron Forton, Sarah MacLachlan, Jenna Parker, Olivia Spagnuolo, Abigail Thiemkey, and Kelsey VandeWetering for their earlier work on this dataset. We thank Dr. Andrew Dennhardt at the Michigan State University Center for Statistical Training and Consulting (MSU CSTAT) for his considerable help with our statistics and model selection. We thank Dr. Alison Ashbury at the Max Planck Institute for Animal Behavior (MPI-AB) for her helpful comments on the manuscript. We thank the Kenyan National Commission for Science, Technology, and Innovation, the Kenya Wildlife Service, the Kenya Wildlife Research and Training Institute, the Narok County Government, the Naboisho Conservancy, and the Mara Conservancy for permission to conduct this research.

## AUTHOR CONTRIBUTIONS

TMM, KDSL, and KEH conceived of the project. KEH and many Mara Hyena Project members (including TMM and KDSL) collected the data. TMM and KDSL supervised SG, KK, and LEM in extracting the data from the field notes. TMM and KDSL processed the data, and TMM performed the statistical analyses, created the visualizations, and wrote the original draft. JCB provided critical insight during data analysis and also during the writing stage of this project. All authors reviewed and edited the manuscript.

## FUNDING

This work was supported by National Science Foundation (NSF) grants OISE-1853934, IOS-1755089, and DEB-1353110 to KEH and JCB, by a National Institutes of Health (NIH) grant R01-GM105042 to KEH, by a Human Frontiers in Science Program (HFSP) grant RGP0051/2019 to KEH, by NSF graduate research fellowships to TMM and KDSL, and by a Michigan State University Dr. Marvin Hensley Research Grant to TMM. Open Access publishing was supported by the Max Planck Digital Library.

## COMPETING INTERESTS

The authors declare no competing interests.

## EXTENDED DATA

**Figure ED1.**
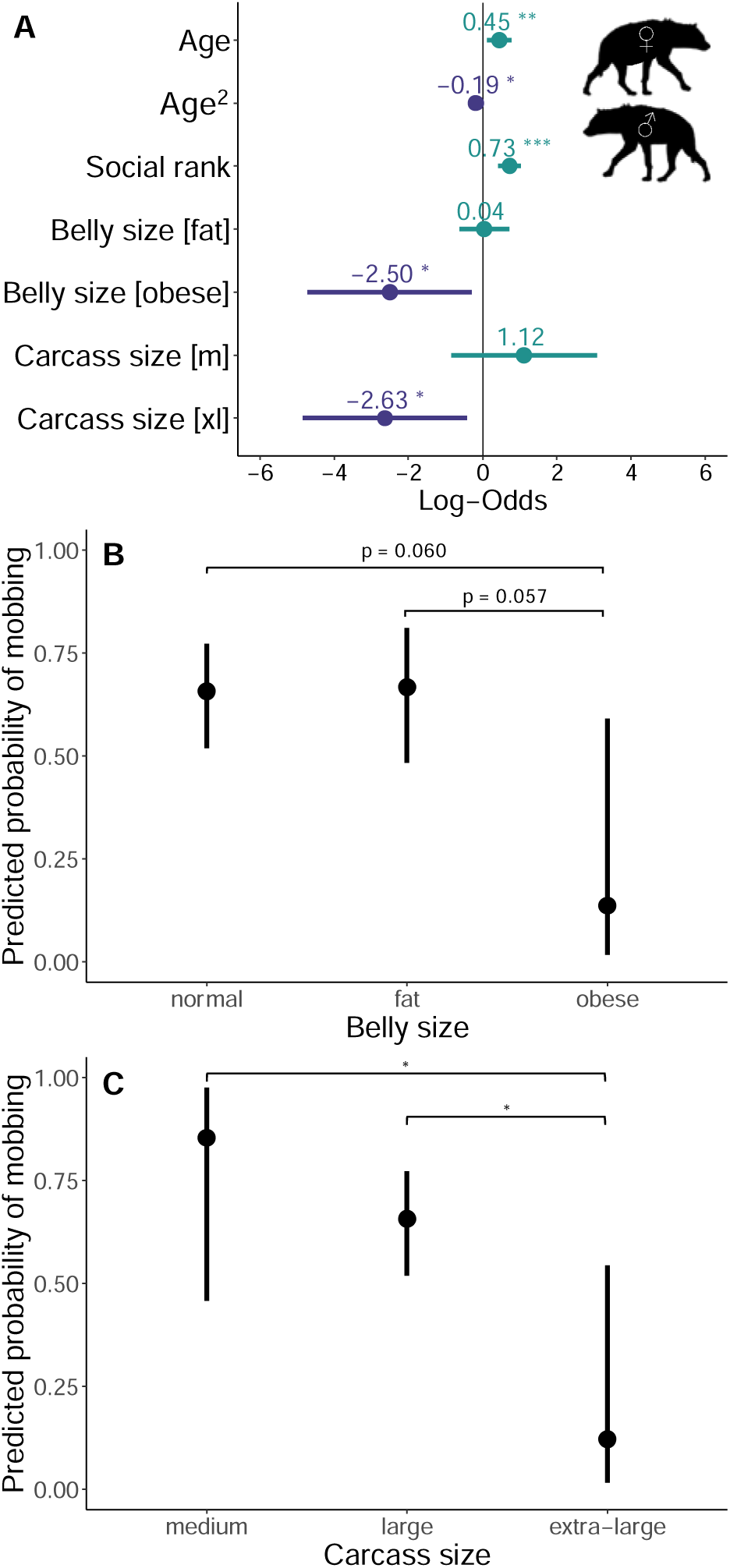
Top model for probability of mobbing participation for adult hyenas in sessions with food (Model I: n-focal hyenas = 407; n-sessions = 34; n-unique hyenas = 195). **A.** Dots depict coefficient estimates, lines depict 95% confidence intervals, and asterisks depict significance at the following p-values: * = 0.05; ** = 0.01; *** = 0.001. **B-C.** Dots depict estimated marginal means and vertical lines depict 95% confidence intervals. Asterisks depict significance in a Tukey post-hoc test at the following p-values: * = 0.05; ** = 0.01; *** = 0.001.

**Figure ED2.**
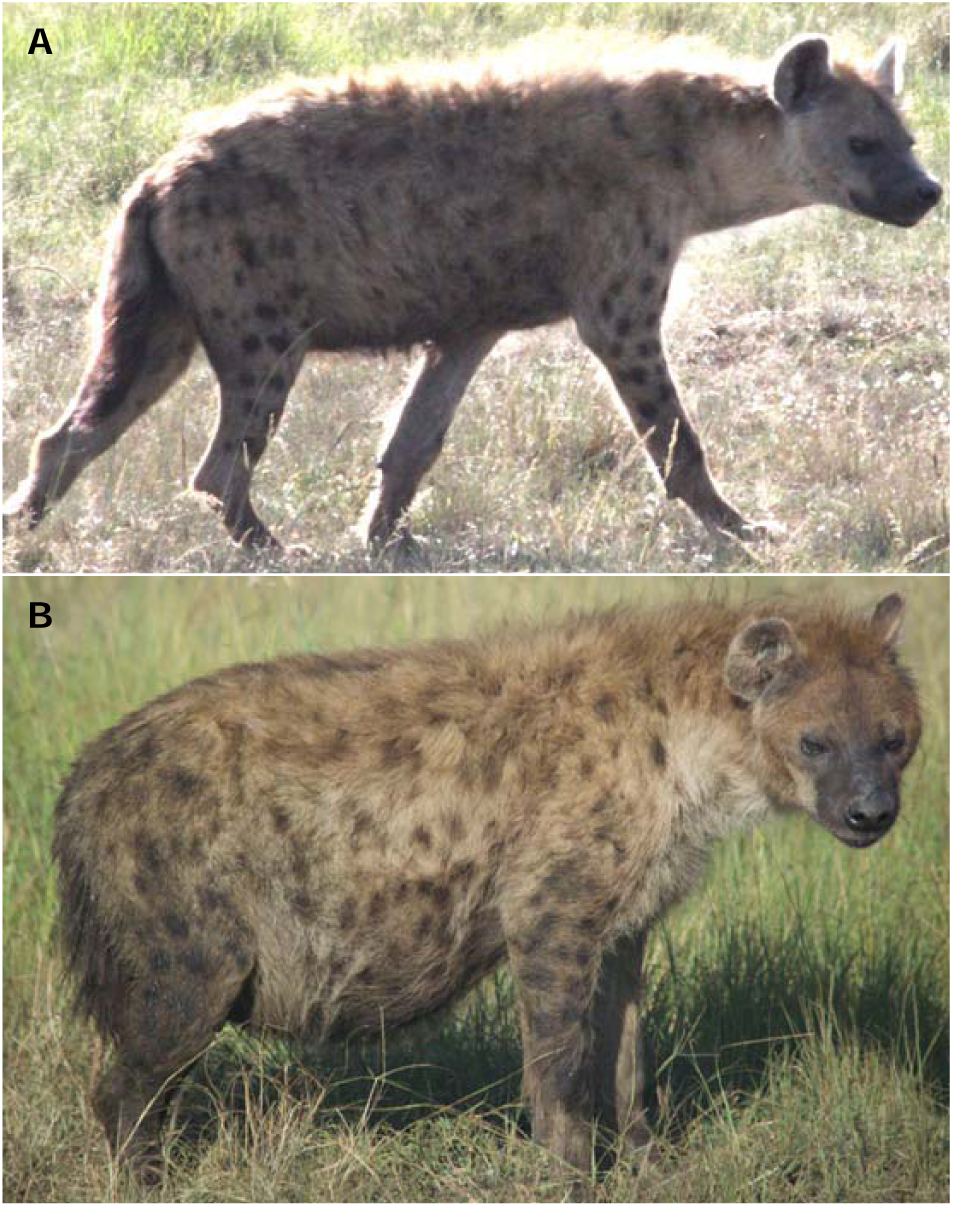
Emma, an adult female hyena with **A.** “normal” belly size and **B.** “obese” belly size. Photos by Oliva S. Spagnuolo.

## Notes

### Competing Interest Statement

The authors have declared no competing interest.

### Summary of Updates

add line numbers, fix figure rendering issue

## REFERENCES

1. Olson, M. The Logic of Collective Action: Public Goods and the Theory of Groups. (Harvard University Press, 1965).

2. Nunn, C. L. & Lewis, R. J. Cooperation and collective action in animal behaviour. in Economics in Nature (eds. Noë, R., Van Hooff, J. A. R. A. M. & Hammerstein, P.) 42–66 (Cambridge University Press, 2001).

3. Dugatkin, L. A. Cooperation Among Animals: An Evolutionary Perspective. (Oxford University Press, 1997).

4. Willems, E. P., Arseneau, T. J. M., Schleuning, X. & van Schaik, C. P. Communal range defence in primates as a public goods dilemma. Philos. Trans. R. Soc. B 370, 20150003 (2015).

5. Kitchen, D. M. & Beehner, J. C. Factors affecting individual participation in group-level aggression among non-human primates. Behaviour 144, 1551–1581 (2007).

6. Hamilton, W. D. The genetical evolution of social behaviour. I. J. Theor. Biol. 7, 1–16 (1964).

7. Zahavi, A. Altruism as a handicap: The limitations of kin selection and reciprocity. J. Avian Biol. 26, 1–3 (1995).

8. Dugatkin, L. A. & Godin, J.-G. J. Prey approaching predators: a cost-benefit perspective. Ann. Zool. Fennici 29, 233–252 (1992).

9. Caro, T. Mobbing and group defense. in Antipredator Defenses in Birds and Mammals 381–412 (University of Chicago Press, 2005).

10. Nunn, C. L. & Deaner, R. O. Patterns of participation and free riding in territorial conflicts among ringtailed lemurs (Lemur catta). Behav. Ecol. Sociobiol. 57, 50–61 (2004).

11. Sumpter, D. J. T. Collective Animal Behavior. (Princeton University Press, 2010).

12. Jolles, J. W., King, A. J. & Killen, S. S. The role of individual heterogeneity in collective animal behaviour. Trends Ecol. Evol. 35, 278–291 (2020).

13. Su, Q., Li, A. & Wang, L. Evolution of cooperation with interactive identity and diversity. J. Theor. Biol. 442, 149–157 (2018).

14. Gavrilets, S. Collective action problem in heterogeneous groups. Philos. Trans. R. Soc. B Biol. Sci. 370, 20150016 (2015).

15. Gavrilets, S. & Fortunato, L. A solution to the collective action problem in between-group conflict with within-group inequality. Nat. Commun. 5, 3526 (2014).

16. Isakov, A., Holcomb, A., Glowacki, L. & Christakis, N. A. Modeling the role of networks and individual differences in inter-group violence. PLoS One 11, 1–10 (2016).

17. Gokcekus, S., Cole, E. F., Sheldon, B. C. & Firth, J. A. Exploring the causes and consequences of cooperative behaviour in wild animal populations using a social network approach. Biol. Rev. 96, 2355–2372 (2021).

18. Holekamp, K. E., Dantzer, B., Stricker, G., Yoshida, K. C. S. & Benson-Amram, S. Brains, brawn and sociality: A hyaena’s tale. Anim. Behav. 103, 237–248 (2015).

19. Kruuk, H. The Spotted Hyena: A Study of Predation and Social Behavior. (University of Chicago Press, 1972).

20. Smith, J. E., Kolowski, J. M., Graham, K. E., Dawes, S. E. & Holekamp, K. E. Social and ecological determinants of fission–fusion dynamics in the spotted hyaena. Anim. Behav. 76, 619–636 (2008).

21. Frank, L. G. Social organization of the spotted hyena (Crocuta crocuta). I. Demography. Anim. Behav. 34, 1500–9 (1986).

22. Van Horn, R. C., Engh, A. L., Scribner, K. T., Funk, S. M. & Holekamp, K. E. Behavioural structuring of relatedness in the spotted hyena (Crocuta crocuta) suggests direct fitness benefits of clan-level cooperation. Mol. Ecol. 13, 449–458 (2004).

23. Frank, L. G. Social organization of the spotted hyaena Crocuta crocuta. II. Dominance and reproduction. Anim. Behav. 34, 1510–1527 (1986).

24. Holekamp, K. E., Smith, J. E., Strelioff, C. C., Van Horn, R. C. & Watts, H. E. Society, demography and genetic structure in the spotted hyena. Mol. Ecol. 21, 613–632 (2012).

25. Smith, J. E., Swanson, E. M., Reed, D. & Holekamp, K. E. Evolution of Cooperation among Mammalian Carnivores and Its Relevance to Hominin Evolution. Curr. Anthropol. 53, S436–S452 (2012).

26. Curio, E. The adaptive significance of avian mobbing: I. Teleonomic hypotheses and predictions. Z. Tierpsychol. 48, 175–183 (1978).

27. Périquet, S., Fritz, H. & Revilla, E. The Lion King and the Hyaena Queen: Large carnivore interactions and coexistence. Biol. Rev. 90, 1197–1214 (2015).

28. Lehmann, K. D. S. et al. Lions, hyenas and mobs (Oh my!). Curr. Zool. 63, 313–322 (2017).

29. Watts, H. E. & Holekamp, K. E. Ecological determinants of survival and reproduction in the spotted hyena. J. Mammal. 90, 461–471 (2009).

30. Crofoot, M. C. Why mob? Reassessing the costs and benefits of primate predator harassment. Folia Primatol. 83, 252–273 (2013).

31. Rusch, H. & Gavrilets, S. The logic of animal intergroup conflict: A review. J. Econ. Behav. Organ. 178, 1014–1030 (2020).

32. Périquet, S. et al. Dynamic interactions between apex predators reveal contrasting seasonal attraction patterns. Oecologia (2021) doi:10.1007/s00442-020-04802-w.

33. Cooper, S. M. Optimal hunting group size: the need for lions to defend their kills against loss to spotted hyaenas. Afr. J. Ecol. 29, 130–6 (1991).

34. Höner, O. P., Wachter, B., East, M. L. & Hofer, H. The response of spotted hyaenas to long-term changes in prey populations: Functional response and interspecific kleptoparasitism. J. Anim. Ecol. 71, 236–246 (2002).

35. Glowacki, L. & Lew-Levy, S. How small-scale societies achieve large-scale cooperation. Curr. Opin. Psychol. 44, 44–48 (2022).

36. Whitham, J. C. & Maestripieri, D. Primate rituals: The function of greetings between male guinea baboons. Ethology 109, 847–859 (2003).

37. DoLJan, G., Glowacki, L. & Rusch, H. Spoils division rules shape aggression between natural groups. Nat. Hum. Behav. 2, 322–326 (2018).

38. Abolins-Abols, M. & Ketterson, E. D. Condition explains individual variation in mobbing behavior. Ethology 123, 495–502 (2017).

39. Samuni, L., Crockford, C. & Wittig, R. M. Group-level cooperation in chimpanzees is shaped by strong social ties. Nat. Commun. 12, 1–10 (2021).

40. Glowacki, L. et al. Formation of raiding parties for intergroup violence is mediated by social network structure. Proc. Natl. Acad. Sci. 113, 12114–12119 (2016).

41. Smith, J. E. et al. Greetings promote cooperation and reinforce social bonds among spotted hyaenas. Anim. Behav. 81, 401–415 (2011).

42. Estes, R. D. & Goddard, J. Prey selection and hunting behavior of the African wild dog. J. Wildl. Manage. 31, 52–70 (1967).

43. Amorós, M., Gil-Sánchez, J. M., López-Pastor, B. de las N. & Moleón, M. Hyaenas and lions: how the largest African carnivores interact at carcasses. Oikos 129, 1820– 1832 (2020).

44. Krause, J., Bumann, D. & Todt, D. Relationship between the position preference and nutritional state of individuals in schools of juvenile roach (Rutilus rutilus). Behav. Ecol. Sociobiol. 30, 177–180 (1992).

45. Glickman, S. E., Frank, L. G., Pavgi, S. & Licht, P. Hormonal correlates of ‘masculinization’ in female spotted hyaenas (Crocuta crocuta). 1. Infancy to sexual maturity. J. Reprod. Fertil. 95, 451–462 (1992).

46. Turner, J. W., Bills, P. S. & Holekamp, K. E. Ontogenetic change in determinants of social network position in the spotted hyena. Behav. Ecol. Sociobiol. 72, 10 (2018).

47. Vullioud, C. et al. Social support drives female dominance in the spotted hyaena. Nat. Ecol. Evol. 3, 71–76 (2019).

48. Watts, H. E., Tanner, J. B., Lundrigan, B. L. & Holekamp, K. E. Post-weaning maternal effects and the evolution of female dominance in the spotted hyena. Proc. R. Soc. B 276, 2291–2298 (2009).

49. Engh, A. L., Esch, K., Smale, L. & Holekamp, K. E. Mechanisms of maternal rank ‘inheritance’ in the spotted hyaena, Crocuta crocuta. Anim. Behav. 60, 323–332 (2000).

50. Jones, S. C., Strauss, E. D. & Holekamp, K. E. Ecology of African Carrion. in Carrion Ecology, Evolution, and Their Applications (eds. Benbow, M. E., Tomberlin, J. K. & Tarone, A. M.) 461–491 (Taylor & Francis Group, 2016).

51. Bergman, T. J. & Beehner, J. C. Measuring social complexity. Anim. Behav. 103, 203–209 (2015).

52. Carter, G. G. et al. Development of New Food-Sharing Relationships in Vampire Bats. Curr. Biol. 30, 1–5 (2020).

53. Whitman, K. L. & Packer, C. A hunter’s guide to aging lions in eastern and southern Africa. (Safari Press & Sports Afield, 2006).

54. Altmann, J. Observational study of behavior: Sampling methods. Behaviour 49, 227– 265 (1974).

55. Green, D. S., Farr, M. T., Holekamp, K. E., Strauss, E. D. & Zipkin, E. F. Can hyena behaviour provide information on population trends of sympatric carnivores? Philos. Trans. R. Soc. B 374, 20180052 (2019).

56. Holekamp, K. E. et al. Patterns of association among female spotted hyenas (Crocuta crocuta). J. Mammal. 78, 55–64 (1997).

57. Farine, D. R. asnipe: Animal Social Network Inference and Permutations for Ecologists. R Packag. version 1.1.12 (2019).

58. Boydston, E. E., Kapheim, K. M., Szykman, M. & Holekamp, K. E. Individual variation in space use by female spotted hyenas. J. Mammal. 84, 1006–1018 (2003).

59. Holekamp, K. E., Szykman, M., Boydston, E. E. & Smale, L. Association of seasonal reproductive patterns with changing food availability in an equatorial carnivore, the spotted hyaena (Crocuta crocuta). J. Reprod. Fertil. 116, 87–93 (1999).

60. Kingdon, J. The Kingdon Field Guide to African Mammals. (Princeton University Press, 2015).

61. Holekamp, K. E., Smale, L. & Szykman, M. Rank and reproduction in the female spotted hyaena. J. Reprod. Fertil. 108, 229–237 (1996).

62. Van Horn, R. C., McElhinny, T. L. & Holekamp, K. E. Age estimation and dispersal in the spotted hyena (Crocuta crocuta). J. Mammal. 84, 1019–1030 (2003).

63. Frank, L. G., Glickman, S. E. & Powch, I. Sexual dimorphism in the spotted hyaena (Crocuta crocuta). J. Zool. London 221, 308–313 (1990).

64. Strauss, E. D. & Holekamp, K. E. Inferring longitudinal hierarchies: Framework and methods for studying the dynamics of dominance. J. Anim. Ecol. 88, 521–536 (2019).

65. Strauss, E. D. DynaRankR: Inferring Longitudinal Dominance Hierarchies. R Packag. version 1.1.0 (2020).

66. East, M. L. & Hofer, H. Male spotted hyenas (Crocuta crocuta) queue for status in social groups dominated by females. Behav. Ecol. 12, 558–568 (2001).

67. Frank, L. G. & Glickman, S. E. Giving birth through a penile clitoris: parturition and dystocia in the spotted hyaena (Crocuta crocuta). J. Zool. London 234, 659–690 (1994).

68. Engh, A. L. et al. Reproductive skew among males in a female-dominated mammalian society. Behav. Ecol. 13, 193–200 (2002).

69. Holekamp, K. E. & Sisk, C. L. Effects of dispersal status on pituitary and gonadal function in the male spotted hyena. Horm. Behav. 44, 385–394 (2003).

70. Curren, L. J., Weldele, M. L. & Holekamp, K. E. Ejaculate quality in spotted hyenas: Intraspecific variation in relation to life-history traits. J. Mammal. 94, 90–99 (2013).

71. Pangle, W. M. & Holekamp, K. E. Age-related variation in threat-sensitive behavior exhibited by spotted hyenas: Observational and experimental approaches. Behaviour 147, 1009–1033 (2010).

72. Watts, H. E. & Holekamp, K. E. Interspecific competition influences reproduction in spotted hyenas. J. Zool. 276, 402–410 (2008).

73. Smith, J. E., Memenis, S. K. & Holekamp, K. E. Rank-related partner choice in the fission–fusion society of the spotted hyena (Crocuta crocuta). Behav. Ecol. Sociobiol. 61, 753–765 (2007).

74. Zuur, A. F., Ieno, E. N. & Elphick, C. S. A protocol for data exploration to avoid common statistical problems. Methods Ecol. Evol. 1, 3–14 (2010).

75. Harrison, X. A. et al. A brief introduction to mixed effects modelling and multi-model inference in ecology. PeerJ 6, e4794 (2018).

76. Magnusson, A. et al. glmmTMB: Generalized Linear Mixed Models using Template Model Builder. R Packag. version 1.0.1 (2020).

77. Bartoń, K. MuMIn: Multi-Model Inference. R Packag. version 1.43.17 (2020).

78. Lüdecke, D., Makowski, D., Waggoner, P., Patil, I. & Ben-Shachar, M. S. performance: Assessment of Regression Models Performance. R Packag. Version 0.7.0 (2021).

79. Hartig, F. DHARMa: Residual Diagnostics for Hierarchical (Multi-Level / Mixed) Regression Models. R Packag. version 0.3.0 (2020).

80. Hothorn, T. et al. multcomp: Simultaneous Inference in General Parametric Models. R Packag. version 1.4-16 (2021).

81. Lüdecke, D. et al. sjPlot: Data Visualization for Statistics in Social Science. R Packag. version 2.8.7 (2021).

82. Lüdecke, D., Aust, F., Crawley, S. & Ben-Shachar, M. S. ggeffects: Create Tidy Data Frames of Marginal Effects for ‘ggplot’ from Model Outputs. R Packag. version 1.0.2 (2021).

83. Wickham, H. et al. ggplot2: Create Elegant Data Visualisations Using the Grammar of Graphics. R Packag. version 3.3.3 (2020).

