## Supplementary Material for "Determinants of hyena participation in risky collective action"

**SUPPLEMENTARY MATERIALS**

**When does cooperative mobbing occur?**

Because of the strong effect of adult male lion presence, we modified our global model of mobbing occurrence (Model A in Table 1) by separating sessions into those where adult male lions were present (Model B in Table S1) and those where male lions were absent (Model C in Table S1). We used these models to inquire whether the number of lions had a continuous effect as a predictor of mobbing occurrence, but we found no support for this. Our top model of the occurrence of mobbing at sessions with adult male lions (Model B: n = 113 complete cases; Table S4) did not include the term for number of lions or for number of adult male lions, nor did any models within 6AIC of the top model. Similarly, our top model of the occurrence of mobbing at sessions without adult male lions (Model C: n = 212 complete cases; Table S4) did not include the term for number of lions, nor did any models within 6AIC of the top model.

**Who participates in cooperative mobbing?**

In our juvenile participation model (Model G: n = 1153 complete cases; Figure S1; Table S4), focal juveniles that were older were more likely to mob than younger juveniles (β = 1.75, p < 0.001). Sex was included in the top model but was not significant (β-male = -0.36, p = 0.362). A near-significant interaction between age and sex indicated that sex differences in mobbing may emerge early in life (β-age x sex = -0.71, p = 0.098). Whether or not the hyena greeted was included in the top model but was not significant (β-greeted = 1.17, p = 0.125). Social rank, association index, and maternal relatedness were not included in the top model or any model within 6 AIC of the top model.

Table S1. Description of outcome variables and predictors used in model selection for mobbing occurrence. (Bolded terms remain in the top model.)

| **Outcome variable** | | **Main effects** | | **Definition*** | **Interaction effects** | | **Random effects** | |
| --- | --- | --- | --- | --- | --- | --- | --- | --- |
| **A. Mobbing occurrence: Logistic model** | | | | | | | | |
| (T/F): Whether a mob occurred during the session | | Session length | | Length of observation session in minutes | Session length x  Number of hyenas present  Session length x  Number of greeters  Number of hyenas present x  Number of lions present  Number of hyenas present x  Male lions present  **Number of greeters x  Number of hyenas present**  Number of greeters x  Number of lions present  **Number of greeters x  Male lions present**  Association index x  Number of hyenas present  Association index x  Number of lions present  Association index x  Male lions present | | None | |
|  |  | Session context  (food, den, other) | | Food (a kill or carcass present), den (an active hyena den present), other (all other sessions) |  |  |  |  |
|  |  | **Prey density** | | **Number of standard deviations from the annual mean prey density** |  |  |  |  |
|  |  | **Number of hyenas present** | | **Total number of hyenas present at the session** |  |  |  |  |
|  |  | Number of lions present | | Total number of lions present at the session |  |  |  |  |
|  |  | **Male lions present (T/F)** | | **(T/F): whether adult male lions were present at the session** |  |  |  |  |
|  |  | **Number of hyenas who greet (greeters)** | | **Number of hyenas who engage in greeting behavior during the session** |  |  |  |  |
|  |  | Mean association index | | Mean association index of all dyads present based on association data from the previous 365 days |  |  |  |  |
| **Outcome variable** | | **Main effects** | | **Definition*** | **Interaction effects** | | **Random effects** | |
| **B. Mobbing occurrence at sessions with adult male lions: Logistic model** | | | | | | | | |
| (T/F): Whether a mob occurred during the session | Session length | | Length of observation session in minutes | | | Session length x  Number of hyenas present  Session length x  Number of greeters  Number of hyenas present x  Number of lions present  Number of hyenas present x  Number of male lions present  Number of greeters x  Number of hyenas present  Number of greeters x  Number of lions present  Number of greeters x  Number of male lions present  Association index x  Number of hyenas present  Association index x  Number of lions present  Association index x  Number of male lions present | | None |
|  | Session context  (food, den, other) | | Food (a kill or carcass present), den (an active hyena den present), other (all other sessions) | | |  |  |  |
|  | **Prey density** | | **Number of standard deviations from the annual mean prey density** | | |  |  |  |
|  | **Number of hyenas present** | | **Total number of hyenas present at the session** | | |  |  |  |
|  | Number of lions present | | Total number of lions present at the session | | |  |  |  |
|  | Number of male lions present | | Total number of adult male lions present at the session | | |  |  |  |
|  | **Number of hyenas who greet (greeters)** | | **Number of hyenas who engage in greeting behavior during the session** | | |  |  |  |
|  | Mean association index | | Mean association index of all dyads present based on association data from the previous 365 days | | |  |  |  |
| **C. Mobbing occurrence at sessions without adult male lions: Logistic model** | | | | | | | | |
| (T/F): Whether a mob occurred during the session | | Session length | | Length of observation session in minutes | Session length x  Number of hyenas present  Session length x  Number of greeters  Number of hyenas present x  Number of lions present  **Number of greeters x  Number of hyenas present**  Number of greeters x  Number of lions present  Association index x  Number of hyenas present  Association index x  Number of lions present | | None | |
|  | | Session context  (food, den, other) | | Food (a kill or carcass present), den (an active hyena den present), other (all other sessions) |  |  |  |  |
|  | | Prey density | | Number of standard deviations from the annual mean prey density |  |  |  |  |
|  | | **Number of hyenas present** | | **Total number of hyenas present at the session** |  |  |  |  |
|  | | Number of lions present | | Total number of lions present at the session |  |  |  |  |
|  | | **Number of hyenas who greet (greeters)** | | **Number of hyenas who engage in greeting behavior during the session** |  |  |  |  |
|  | | Mean association index | | Mean association index of all dyads present based on association data from the previous 365 days |  |  |  |  |
| *See Model Predictors section in Methods for a detailed description of these variables. | | | | | | | | |

Table S2. Description of outcome variables and predictors used in model selection for mobbing participation. (Bolded terms remain in the top model.)

| **Outcome variable** | **Main effects** | | **Definition*** | | | **Interaction effects** | | **Random effects** |
| --- | --- | --- | --- | --- | --- | --- | --- | --- |
| **D. Mobbing participation by all hyenas: Logistic mixed-effects model** | | | | | | | | |
| (T/F): Whether a hyena participated in the mob | | **Age** | | **Age of focal hyena in years** | None | | Session ID-Mob ID  Hyena ID | |
|  |  | **Sex** | | **Sex of focal hyena** |  | |  |  |
| **E. Mobbing participation by adult female hyenas: Logistic mixed-effects model** | | | | | | | | |
| (T/F): Whether a hyena participated in the mob | | **Age** | | **Age of focal hyena in years** | Age x Social rank  **Social rank x Greeted**  **Social rank x  Association index**  Social rank x  Maternal relatedness | | Session ID-Mob ID  Hyena ID | |
|  |  | **Social rank** | | **Social rank of focal hyena during calendar year of session** |  |  |  |  |
|  |  | Reproductive state (nulliparous, pregnant, lactating, other) | | Nulliparous (never given birth), pregnant (pregnant with at least one cub), lactating (nursing at least one cub), other (cycling or between pregnancies) |  |  |  |  |
|  |  | **Greeted (T/F)** | | **(T/F): whether focal hyena engaged in greeting behavior in five minutes prior to mob** |  |  |  |  |
|  |  | **Association index with participants** | | **Mean association index between focal hyena and participants based on association data from the previous 365 days** |  |  |  |  |
|  |  | **Maternal relatedness with participants** | | **Proportion of participants to whom focal hyena is closely maternally related** |  |  |  |  |
| **F. Mobbing participation by adult male hyenas: Logistic mixed-effects model** | | | | | | | | |
| (T/F): Whether a hyena participated in the mob | | **Age** | | **Age of focal hyena in years** | Social rank x Greeted  Social rank x  Association index | | Session ID-Mob ID  Hyena ID | |
|  |  | **Social rank** | | **Social rank of focal hyena during calendar year of session** |  |  |  |  |
|  |  | Dispersal status (natal, immigrant) | | Natal (born in clan), immigrant (immigrated into clan) |  |  |  |  |
|  |  | Greeted (T/F) | | (T/F): whether focal hyena engaged in greeting behavior in five minutes prior to mob |  |  |  |  |
|  |  | **Association index with participants** | | **Mean association index between focal hyena and participants based on association data from the previous 365 days** |  |  |  |  |
| **Outcome variable** | | **Main effects** | | **Definition*** | **Interaction effects** | | **Random effects** | |
| **G. Mobbing participation by juvenile hyenas: Logistic mixed-effects model** | | | | | | | | |
| (T/F): Whether a hyena participated in the mob | | **Age** | | **Age of focal hyena in years** | **Age x Sex**  Age x Social rank  Sex x Social rank | | Session ID-Mob ID  Hyena ID | |
|  |  | **Sex** | | **Sex of focal hyena** |  |  |  |  |
|  |  | Social rank | | Social rank of focal hyena during calendar year of session |  |  |  |  |
|  |  | **Greeted (T/F)** | | **(T/F): whether focal hyena engaged in greeting behavior in five minutes prior to mob** |  |  |  |  |
|  |  | Association index with participants | | Mean association index between focal hyena and participants based on association data from the previous 365 days |  |  |  |  |
|  |  | Maternal relatedness with participants | | Proportion of participants to whom focal hyena is closely maternally related |  |  |  |  |
| *See Model Predictors section in Methods for a detailed description of these variables. | | | | | | | | |

Table S3. Description of outcome variables and predictors used in model selection for potential individual benefits of mobbing. (Bolded terms remain in the top model.)

| **Outcome variable** | **Main effects** | **Definition*** | **Interaction effects** | **Random effects** |
| --- | --- | --- | --- | --- |
| **H. Are mobs more likely to occur at sessions with higher quality and/or larger food? Logistic model** | | | | |
| (T/F): Whether a mob occurred during the session | Session length | Length of observation session in minutes | Session length x  Number of hyenas present  Session length x  Number of greeters  Number of hyenas present x  Number of lions present  Number of hyenas present x  Male lions present  Number of greeters x  Number of hyenas present  Number of greeters x  Number of lions present  **Number of greeters x  Male lions present**  Association index x  Number of hyenas present  Association index x  Number of lions present  Association index x  Male lions present | None |
|  | Prey density | Number of standard deviations from the annual mean prey density |  |  |
|  | **Number of hyenas present** | **Total number of hyenas present at the session** |  |  |
|  | Number of lions present | Total number of lions present at the session |  |  |
|  | **Male lions present (T/F)** | **(T/F): whether adult male lions were present at the session** |  |  |
|  | **Number of hyenas who greet (greeters)** | **Number of hyenas who engage in greeting behavior during the session** |  |  |
|  | Mean association index | Mean association index of all dyads present based on association data from the previous 365 days |  |  |
|  | **Carcass freshness (fresh, old)** | **Fresh (the prey was recently killed), old (the prey was killed >24h ago)** |  |  |
|  | Carcass size (medium, large, extra-large) | Medium (<20-100 kg), large (100-500 kg), extra-large (>500 kg) |  |  |
| **I. Are hyenas in poorer nutritional condition more likely to mob at sessions with food? Logistic mixed-effects model** | | | | |
| (T/F): Whether a hyena mobbed during the session | **Age** | **Age of focal hyena in years** | Social rank x Belly size Social rank x Carcass size | Session ID  Hyena ID |
|  | Sex | Sex of focal hyena |  |  |
|  | **Social rank** | **Social rank of focal hyena during calendar year of session** |  |  |
|  | **Belly size (normal, fat, obese)** | **Normal (fit but not fat), fat (big full belly), obese (monstrously giant belly)** |  |  |
|  | Carcass freshness (fresh, old) | Fresh (the prey was recently killed), old (the prey was killed >24h ago) |  |  |
|  | **Carcass size (medium, large, extra-large)** | **Medium (< 20-100 kg), large (100-500 kg), extra-large (> 500 kg)** |  |  |
| **Outcome variable** | **Main effects** | **Definition*** | **Interaction effects** | **Random effects** |
| **J. Are hyenas who mob more likely to feed immediately after the mob? Logistic mixed-effects model** | | | | |
| (T/F): Whether a hyena fed in the five minutes after the mob | **Age** | **Age of focal hyena in years** | Social rank x Participant  Participant x Carcass freshness  Participant x Carcass size | Session ID**  Hyena ID |
|  | Sex | Sex of focal hyena |  |  |
|  | **Social rank** | **Social rank of focal hyena during calendar year of session** |  |  |
|  | **Participant (T/F)** | **(T/F): whether focal hyena participated in the mob** |  |  |
|  | Carcass freshness (fresh, old) | Fresh (the prey was recently killed), old (the prey was killed >24h ago) |  |  |
|  | Carcass size (medium, large, extra-large) | Medium (< 20-100 kg), large (100-500 kg), extra-large (> 500 kg) |  |  |
| **K. Are hyenas who mob more likely to feed later during the session? Logistic mixed-effects model** | | | | |
| (T/F): Whether a hyena fed during the session | Age | Age of focal hyena in years | Social rank x Participant  Participant x Carcass size | Session ID*** |
|  | **Sex** | **Sex of focal hyena** |  |  |
|  | **Social rank** | **Social rank of focal hyena during calendar year of session** |  |  |
|  | Participant (T/F) | (T/F): whether focal hyena mobbed during the session |  |  |
|  | Carcass freshness (fresh, old) | Fresh (the prey was recently killed), old (the prey was killed >24h ago) |  |  |
|  | Carcass size (medium, large, extra-large) | Medium (< 20-100 kg), large (100-500 kg), extra-large (> 500 kg) |  |  |
| *See Model Predictors section in Methods for a detailed description of these variables.  **The random effect of Mob ID was removed from this global model because it explained no variance.  ***The random effect of Hyena ID was removed from this global model because it explained no variance. | | | | |

**Table S4.** All top logistic mixed-effects models presented in the manuscript.

| **Predictors** | **Log-Odds** | **SE** | **p** |
| --- | --- | --- | --- |
| **A. Mobbing occurrence (n-obs = 321)** | | | |
| Number of hyenas present | 0.87 | 0.17 | **<0.001** |
| Male lions present [TRUE] | -0.73 | 0.30 | **0.014** |
| Prey density | 0.27 | 0.13 | **0.038** |
| Number of hyenas who greet (greeters) | 0.53 | 0.22 | **0.017** |
| Number of hyenas present x Number of greeters | -0.32 | 0.17 | 0.061 |
| Male lions present x Number of greeters | 0.67 | 0.35 | 0.059 |
| *Marginal R^2^ = 0.261* |  |  |  |
| **B. Mobbing occurrence at sessions with adult male lions (n-obs = 113)** | | | |
| Number of hyenas present | 1.26 | 0.35 | **<0.001** |
| Prey density | 0.63 | 0.25 | **0.010** |
| Number of hyenas who greet (greeters) | 0.93 | 0.32 | **0.004** |
| *Marginal R^2^ = 0.371* |  |  |  |
| **C. Mobbing occurrence at sessions without adult male lions (n-obs = 212)** | | | |
| Number of hyenas present | 0.69 | 0.19 | **<0.001** |
| Number of hyenas who greet (greeters) | 0.66 | 0.26 | **0.012** |
| Number of hyenas present x Number of greeters | -0.41 | 0.20 | **0.047** |
| *Marginal R^2^ = 0.188* |  |  |  |
| **D. Mobbing participation by all hyenas   (n-obs = 4383, n-sessions = 117, n-mobs = 342, n-hyenas = 431)** | | | |
| Age | 0.72 | 0.09 | **<0.001** |
| Age^2^ | -0.40 | 0.05 | **<0.001** |
| Sex [male] | -1.04 | 0.16 | **<0.001** |
| *ICC =* 0.52 |  |  |  |
| *Marginal R^2^ / Conditional R^2^ =* 0.090 / 0.566 |  |  |  |
| **E. Mobbing participation by adult female hyenas   (n-obs = 2280, n-sessions = 109, n-mobs = 323, n-hyenas = 169)** | | | |
| Age | 0.08 | 0.10 | 0.410 |
| Age^2^ | -0.13 | 0.05 | **0.014** |
| Social rank | 0.17 | 0.10 | 0.113 |
| Greeted [TRUE] | 1.17 | 0.25 | **<0.001** |
| Association index (participants) | 0.39 | 0.13 | **0.004** |
| Maternal relatedness (participants) | 0.23 | 0.09 | **0.013** |
| Greeted x Social rank | -0.75 | 0.29 | **0.009** |
| Association index x Social rank | 0.21 | 0.09 | **0.024** |
| *ICC =* 0.51 |  |  |  |
| *Marginal R^2^ / Conditional R^2^ =* 0.092 / 0.554 |  |  |  |
| **Predictors** | **Log-Odds** | **SE** | **p** |
| **F. Mobbing participation by adult male hyenas   (n-obs = 893, n-sessions = 101, n-mobs = 288, n-hyenas = 124)** | | | |
| Age | 0.12 | 0.25 | 0.616 |
| Age^2^ | -0.37 | 0.17 | **0.025** |
| Social rank | 0.97 | 0.22 | **<0.001** |
| Association index (participants) | 0.36 | 0.18 | **0.045** |
| *ICC =* 0.66 |  |  |  |
| *Marginal R^2^ / Conditional R^2^ =* 0.106 / 0.696 |  |  |  |
| **G. Mobbing participation by juvenile hyenas   (n-obs = 1153, n-sessions = 88, n-mobs = 269, n-hyenas = 247)** | | | |
| Age | 1.75 | 0.33 | **<0.001** |
| Sex [male] | -0.36 | 0.39 | 0.362 |
| Greeted [TRUE] | 1.17 | 0.76 | 0.125 |
| Age x Sex | -0.71 | 0.43 | 0.098 |
| *ICC =* 0.66 |  |  |  |
| *Marginal R^2^ / Conditional R^2^ =* 0.178 / 0.722 |  |  |  |
| **H. Mobbing occurrence at sessions with food (n-obs = 218)** | | | |
| Number of hyenas present | 0.84 | 0.20 | **<0.001** |
| Male lions present [TRUE] | -0.65 | 0.35 | 0.061 |
| Number of hyenas who greet (greeters) | 0.10 | 0.20 | 0.638 |
| Carcass freshness [fresh] | -0.32 | 0.42 | 0.443 |
| Male lions present x Number of greeters | 0.99 | 0.44 | **0.025** |
| *Marginal R^2^ = 0.216* |  |  |  |
| **I. Mobbing participation by hyenas of variable body condition at sessions with food   (n-obs = 407, n-sessions = 34, n-hyenas = 195)** | | | |
| Age | 0.45 | 0.17 | **0.007** |
| Age^2^ | -0.19 | 0.09 | **0.033** |
| Social rank | 0.73 | 0.16 | **<0.001** |
| Belly size [fat] | 0.04 | 0.34 | 0.898 |
| Belly size [obese] | -2.50 | 1.13 | **0.027** |
| Carcass size [medium] | 1.12 | 1.00 | 0.265 |
| Carcass size [extra-large] | -2.63 | 1.13 | **0.020** |
| *ICC =* 0.31 |  |  |  |
| *Marginal R^2^ / Conditional R^2^ =* 0.181 / 0.434 |  |  |  |
| **J. Feeding immediately after the mob   (n-obs = 1049, n-sessions = 41, n-hyenas = 185)** | | | |
| Age | 0.08 | 0.18 | 0.668 |
| Age^2^ | -0.26 | 0.09 | **0.005** |
| Social rank | 0.42 | 0.15 | **0.006** |
| Participant [TRUE] | 0.56 | 0.20 | **0.006** |
| *ICC =* 0.48 |  |  |  |
| *Marginal R^2^ / Conditional R^2^ =* 0.082 / 0.519 |  |  |  |
| **Predictors** | **Log-Odds** | **SE** | **p** |
| **K. Feeding during the session   (n-obs = 673, n-sessions = 59)** | | | |
| Sex [male] | -0.66 | 0.26 | **0.012** |
| Social rank | 0.24 | 0.13 | 0.069 |
| *ICC =* 0.38 |  |  |  |
| *Marginal R^2^ / Conditional R^2^ =* 0.048 / 0.409 |  |  |  |

**Figure S1**. Top model for probability of mobbing participation by juvenile hyenas (Model G: n-focal hyenas = 1153; n-sessions = 88; n-mobs = 269; n-unique hyenas = 247). **A.** Dots depict coefficient estimates, lines depict 95% confidence intervals, and asterisks depict significance at the following p-values: * = 0.05; ** = 0.01; *** = 0.001. **B.** Lines depict estimated marginal means and shaded areas depict 95% confidence intervals.

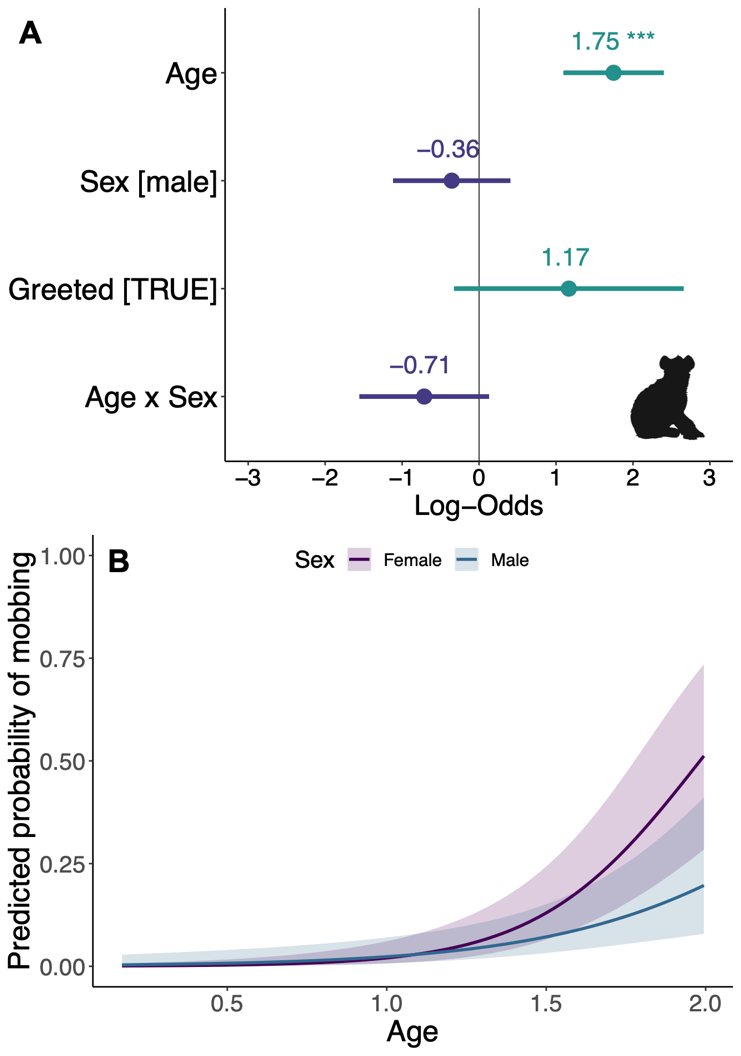

**Table S5.** Food sessions at different carcass sizes.

| **Carcass size** | **Number of sessions** | **Number of sessions with mobbing** |
| --- | --- | --- |
| XL | 15 | 7 |
| L | 131 | 58 |
| M | 35 | 17 |

**Table S6**. Categorization of carcass size based on prey species and prey age.

| **Prey species** | **Carcass size** | |
| --- | --- | --- |
|  | **Juvenile** | **Adult** |
| Buffalo (*Syncerus caffer*), Elephant (*Loxodonta africana*),  Giraffe (*Giraffa camelopardalis*), Hippopotamus (*Hippopotamus amphibius*) | XL | XL |
| Domestic cow (*Bos taurus*), Hartebeest (*Alcelaphus buselaphus*),  Topi (*Damaliscus lunatus*), Wildebeest (*Connochaetes gnou*),  Zebra (*Equus quagga*) | M | L |
| Domestic goat (*Capra aegagrus*), Domestic sheep (*Ovis aries*),  Grant's gazelle (*Nanger granti*), Impala (*Aepyceros melampus*),  Thompson's gazelle (*Eudorcas thomsonii*), Warthog (*Phacochoerus africanus*) | S | M |
